# Cell Growth and Division Shape mRNA–Protein Correlations

**DOI:** 10.64898/2026.05.04.722628

**Authors:** Kuheli Biswas, Michael Sheinman, Leonardo A Sepúlveda, Ido Golding, Ariel Amir

**Affiliations:** Department of Physics of Complex Systems, Weizmann Institute of Science, Rehovot, Israel; Howard Hughes Medical Institute, Harvard University, Cambridge, Massachusetts, USA; Department of Chemistry and Chemical Biology, Harvard University, Cambridge, Massachusetts, USA; Department of Physics, Harvard University, Cambridge, Massachusetts, USA; Department of Physics, University of Illinois at Urbana-Champaign, Urbana, Illinois, USA

**Author notes:** These authors contributed equally (Kuheli Biswas, Michael Sheinman).

## Abstract

Correlations between cellular variables, such as gene-expression levels, provide insights into regulatory mechanisms. We focus here on correlations between mRNA and protein levels and re-examine previously derived analytical predictions. We test this prediction on single-cell *E. coli* data and see substantial disagreement. We hypothesize that this discrepancy arises from the assumption of constant cell volume and develop a theoretical framework for mRNA–protein correlations in growing and dividing cells. Within this framework, we derive an analytical expression for mRNA– protein correlations and show that explicit incorporation of growth and division substantially alters these correlations. The resulting relation is invariant to upstream transcriptional dynamics, and we validate it using stochastic simulations across multiple gene-regulatory architectures. Finally, we show that the derived predictions are consistent with the *E. coli* data.

## 2 Introduction

Gene expression is a fundamental process in cellular physiology, governed by a complex regulatory network of mRNAs and proteins. The resulting network dynamics generate rich correlation structures among mRNA and protein levels, which can reveal features of the underlying gene-regulatory mechanisms. These correlations are further modulated by noise, which may exert opposing effects. In the classical dual-reporter setup of Elowitz *et al*. [1], intrinsic noise decreases the correlation between reporter expression levels. In other network architectures, however, intrinsic noise of an upstream regulator can propagate through a genetic cascade, thereby increasing correlations between downstream expression levels [2, 3, 4]. In these and other related studies [5, 6, 7], specific, relatively small regulatory networks were analyzed, leaving unresolved the structure of correlations and their relationship to intrinsic noise in generic regulatory circuits with many interacting components, feedback loops, and stochastic molecular processes.

To this end, seminal studies have derived exact relations between measurable quantities that depend only on a limited subset of interactions [8, 9, 10]. It was shown that mRNA–protein correlations obey a universal relation that is independent of kinetic schemes of mRNA production and are determined solely by protein production and degradation dynamics [8]. Specifically, when the translation rate is proportional to the mRNA abundance, the correlation between mRNA copy number, *M*, and protein copy number, *P*, is predicted by [8]

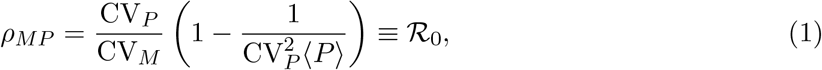

which depends on the mean protein copy number ⟨*P* ⟩, and the intrinsic noise in protein and mRNA copy numbers, respectively, denoted by CV_*P*_ and CV_*M*_.

In this study, we reexamine Eq. (1) and show that it is inconsistent with single-cell *E. coli* data [11]. We hypothesized that this discrepancy arises from the assumption of constant-volume, made in Ref. [8]. This interpretation is supported by earlier studies showing that cell growth and division can substantially alter the statistics of mRNA and protein abundance relative to constant-volume models [12, 13, 14, 15, 16, 17]. Here, we develop a general framework for deriving mRNA–protein correlations for a gene embedded in an arbitrary regulatory network while explicitly accounting for cell growth and division. Within this framework, we find in simulations that Eq. (1) does not hold, and that this discrepancy persists even when the relation is reformulated in terms of concentrations. We then derive an expression for the mRNA–protein correlation in growing and dividing cells and validate this prediction using stochastic simulations across several upstream transcriptional network architectures. Finally, we test our prediction against the same single-cell *E. coli* data [11] and find good agreement with the measurements.

## 3 Results

### 3.1 Comparison of the constant-volume model with experimental data

We begin by comparing Eq. (1), derived from the constant-volume model, with experimental data. To this end, we use single-cell measurements of mRNA and protein copy numbers in *E. coli* across multiple growth conditions and regulatory architectures, previously reported in Ref. [11]. The dataset contains two types of mRNA–protein pairs: *(i)* pairs in which the mRNA and protein were measured for the same gene (hereafter referred to as same-gene pairs), and *(ii)* pairs in which the mRNA and protein were measured for different genes (hereafter referred to as different-gene pairs). For same-gene pairs, measurements are available both in the presence and absence of protein-mediated feedback regulation. The corresponding regulatory architectures are shown in Fig. 1(a). In the experiment [11], protein copy numbers were quantified using immunofluorescence, whereas mRNA copy numbers were measured by single-molecule fluorescence *in situ* hybridization.

**Figure 1:**
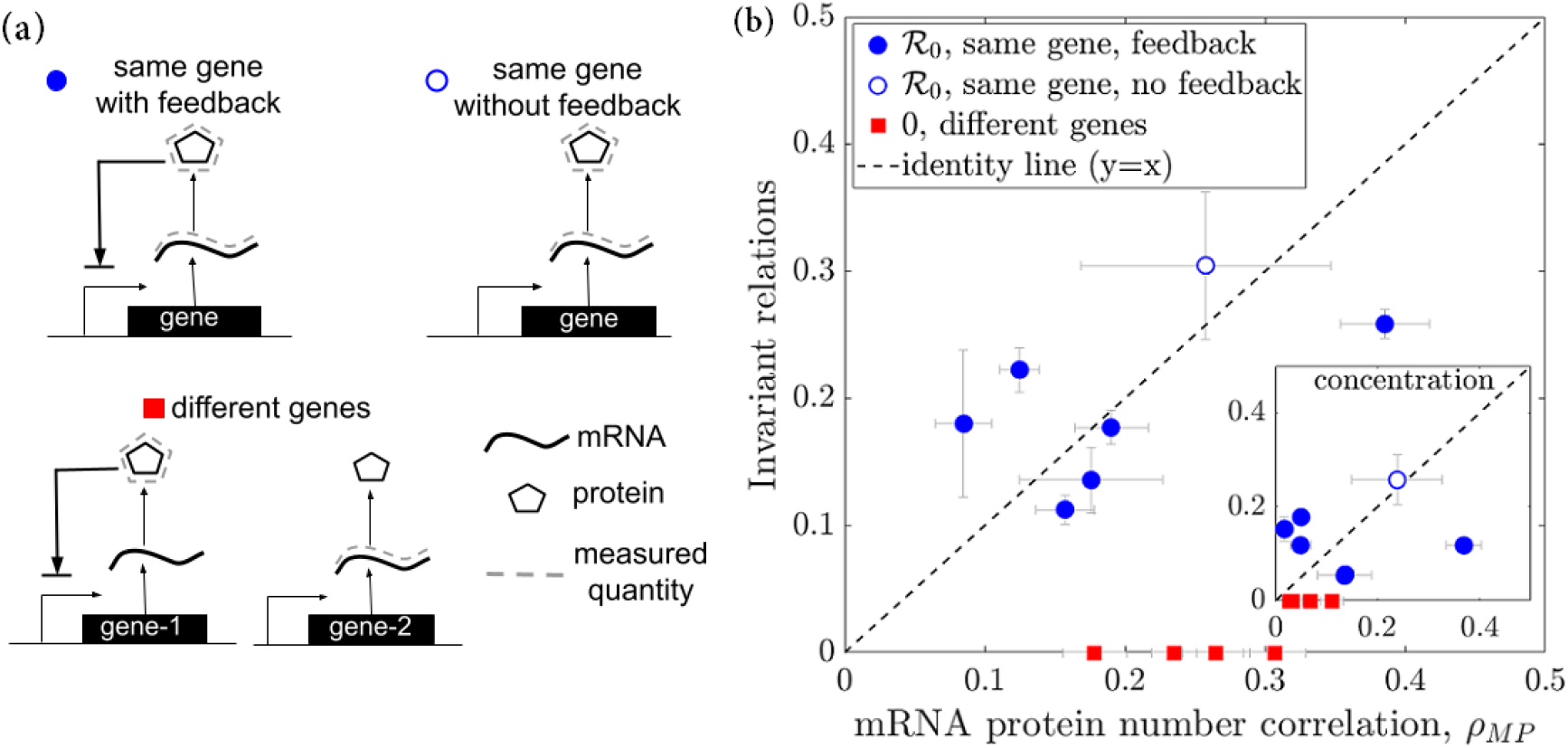
Measured mRNA–protein correlations in *E. coli* deviate from the constantvolume prediction of Eq. (1). **(a)** Schematic of the three classes of mRNA–protein pairs analyzed in the *E. coli* dataset [11]: same-gene pairs with feedback (blue filled circles), samegene pairs without feedback (blue open circles), and different-gene pairs (red filled squares). **(b)** Comparison between measured mRNA–protein copy-number correlations, *ρ*_*MP*_, and the constantvolume prediction ℛ_0_ in Eq. (1) for same-gene pairs (blue circles). Error bars indicate bootstrap standard deviations obtained by resampling individual cells. The dashed line denotes *y* = *x*, corresponding to perfect agreement between prediction and measurement. For different-gene pairs, the constant-volume model predicts zero correlation, provided that the two genes neither regulate one another nor share a common regulator. In contrast, the experimental data exhibit substantial nonzero mRNA–protein correlations for these pairs (red squares). In the inset, we plot the same, but for mRNA and protein concentrations instead of copy numbers.

The constant-volume model developed in Ref. [8] predicts that same-gene mRNA–protein pairs should satisfy Eq. (1), whereas different-gene pairs should exhibit zero correlation, provided that the two genes neither regulate one another nor share a common regulator. However, when tested against the experimental data, for same-gene pairs, Eq. (1) fails in 2 of the 7 growth conditions examined, with Bonferroni-corrected *p*-values of 0.03 and 0.01. Moreover, for different-gene pairs, the constant-volume prediction of zero correlation clearly fails in all four studied growth conditions (all *p*-values are below 10^−7^). A summary of these results is shown in Fig. 1(b). One might argue heuristically that Eq. (1) could be adapted by replacing mRNA and protein abundances with their concentrations. However, as shown in the inset of Fig. 1(b), this concentration-based relation is also inconsistent with the data, both for same-gene pairs (4 growth conditions with Bonferroni-corrected *p <* 10^−3^) and for different-gene pairs (2 growth conditions with *p <* 10^−3^).

Several factors may contribute to these discrepancies, including limited sample size and possible experimental artifacts. Nevertheless, we hypothesize that the dominant source of the disagreement is the assumption of constant cell volume under which Eq. (1) was derived [8]. The remainder of this article is devoted to establishing this hypothesis and analyzing its consequences. In the following section, we derive the mRNA–protein copy-number correlation for a gene embedded in a regulatory network while explicitly accounting for cell growth and division, and compare the resulting prediction with simulations and the experimental data.

### 3.2 Invariant relations for growing and dividing cells

We consider a population in which each cell grows and divides into two daughter cells via binary fission, generating a branching lineage tree [18, 19] (see Fig. 2(a)). Within each cell, we consider the expression of a gene embedded in an arbitrary, unspecified network that affects transcription dynamics, while we specify the translation process (discussed in detail in the following sections). We make no assumptions about the growth process or the division-control mechanism. At division, cellular components are partitioned independently between the two daughter cells according to binomial statistics.

**Figure 2:**
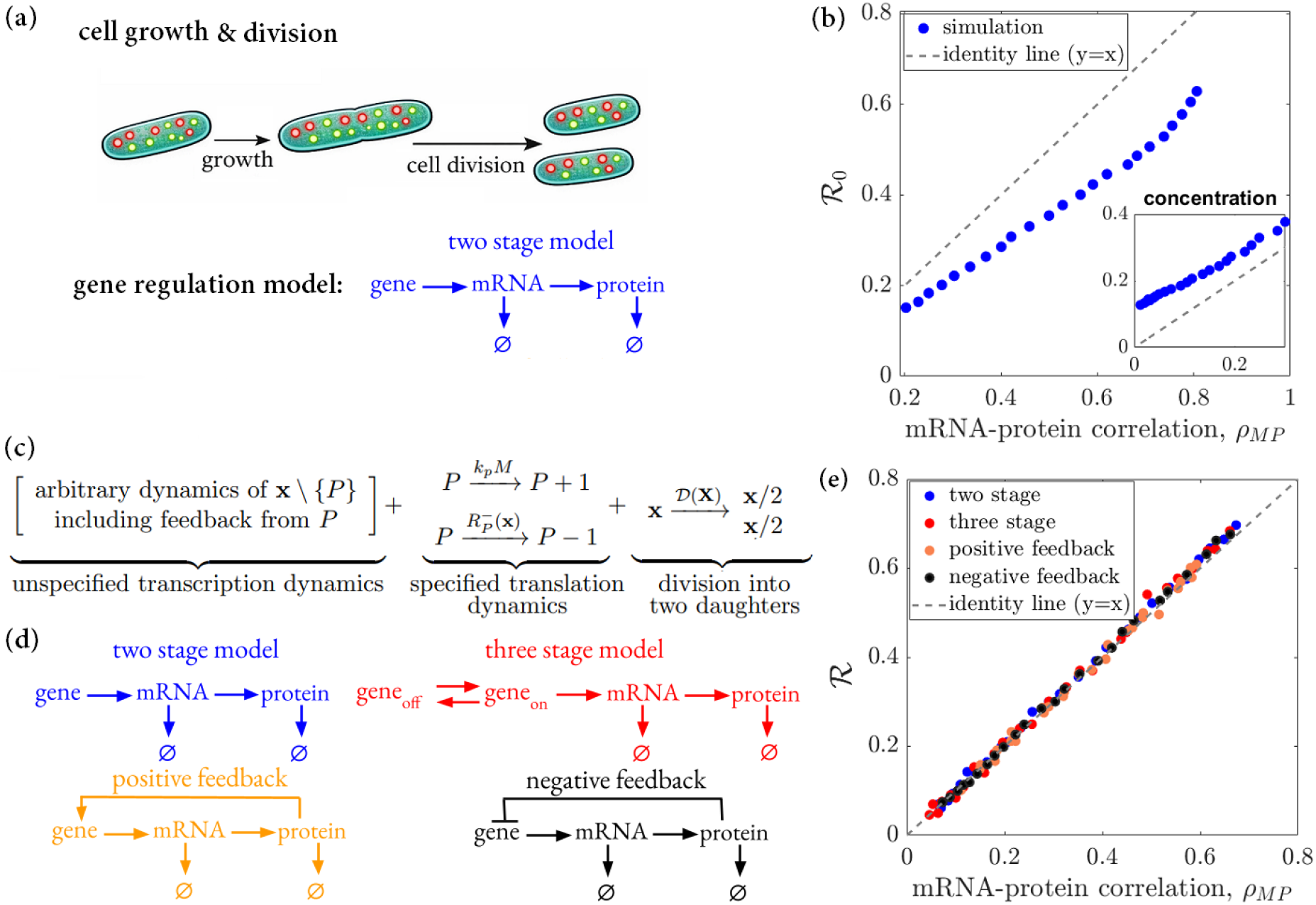
mRNA-protein correlation for growing and dividing cells. **(a)** Schematic of the growing-dividing model. Individual cells grow and divide into two daughter cells, while intracellular mRNA and protein copy numbers (yellow and red dots, respectively) evolve stochastically (top). The underlying gene-expression dynamics are represented by a two-stage regulatory module, in which a gene produces mRNA, which in turn produces protein, with both mRNA and protein undergoing degradation (bottom). **(b)** mRNA–protein correlation, *ρ*_*MP*_, for growing-dividing cells deviates from the invariant prediction of Eq. (1). Inset **(b)**: the corresponding comparison for mRNA–protein concentration correlations, *ρ*[*M* ][*P* ]. **(c)** For arbitrary dynamics of cellular components excluding proteins (**x** \ {*P* }), translation is modeled as a first-order process, with proteins produced at rate *k*_*p*_*M*. Protein degradation is allowed to follow arbitrary kinetics, described by the flux 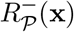, which may depend on the full cellular state. Upon division, all cellular components are partitioned between two daughter cells, with division occurring at a state-dependent rate 𝒟(**x**). **(d)** We consider several upstream transcription networks: a two-stage model (gene always active), a three-stage model (gene switches between active and inactive states at a constant rate) [25], and models with positive or negative feedback [26, 27]. For the two-stage and three-stage models, the mRNA production rate is *k*_*m*_*g* [25]. For positive and negative feedback, the mRNA production rate follows a Hill function [26, 27], 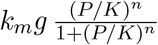 and 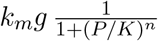, respectively, where *K* is the dissociation constant (half-maximal regulation) and *n* is the Hill coefficient (regulatory steepness). In all cases, the mRNA degradation rate is *γ*_*m*_*M*. Definitions of model parameters are provided in Appendix Table 1. **(e)** Simulation results for *ρ*_*MP*_ across the regulatory networks in (d) collapse onto the invariant prediction *ρ*_*MP*_ =ℛ (Eq. 5), indicating universality with respect to upstream network architecture. In all panels, symbols denote stochastic simulations [28] of 10^3^ cells.

Within this framework, the probability density of the cell state, 𝒫(**x**, *t*) (for brevity, we sometimes omit the explicit time dependence below), is governed by the master equation

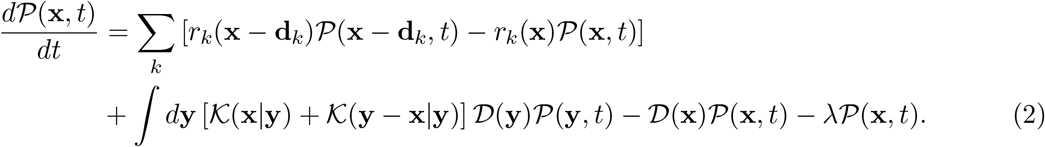

Here, *r*_*k*_(**x** − **d**_*k*_) 𝒫(**x** − **d**_*k*_) is the probability flux from state **x** − **d**_*k*_ to state **x**, and 𝒟(**x**) is the state-dependent division rate. The sum runs over all possible transitions *k*, and each transition is represented by an integer vector **d**_*k*_, which is the difference between two states. The quantity 𝒦(**x**|**y**) denotes the binomial partition kernel, i.e., the conditional probability that a daughter cell is born in state **x** given that the mother cell divided in state **y**. The term −*λ* 𝒫(**x**, *t*) accounts for the exponentially growing population with growth rate *λ*, such that ∫ 𝒫(**x**, *t*) *d***x** = 1. Using the master equation (2), we derive an invariant relation for mRNA–protein copy-number correlations.

Taking the steady state limit of Eq. (2), we obtain a relation for the correlation between the protein copy number *P* and its production rate *Γ* :

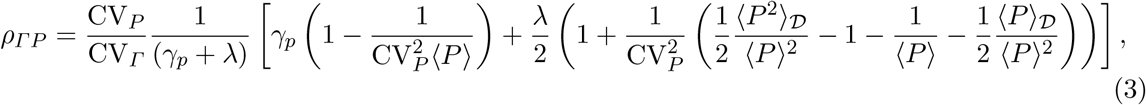

which is derived in Appendix Sec. A. Here *γ*_*p*_ is the protein degradation rate, ⟨*P* ⟩_𝒟_ and ⟨*P* ^2^⟩_𝒟_ are, respectively, the first and second moments of the protein copy number at the time of cell division. Unless stated otherwise, all statistics are computed over the population ensemble. Notably, as in Ref. [8], this result does not depend on assuming first-order protein degradation or any other specific degradation mechanism. Rather, we require only that proteins have a constant mean lifetime, equal to 1*/γ*_*p*_. By Little’s law [20], the mean rate of protein loss then satisfies 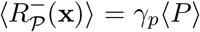, which we use in the Appendix to derive Eq. (3). Eq. (3) also applies to populations with random removal of cells (e.g., chemostats [21] or turbidostats [22]) and to fixed-size-population turnover models such as Moran processes [23]. Furthermore, Eq. (3) is invariant in the sense that its validity does not depend on the specific mechanisms governing mRNA dynamics, protein production and degradation, genome replication, or division control. In the limit where protein degradation is negligible compared with growth-mediated dilution, *γ*_*p*_ ≪ *λ* [24], Eq. (3) is reduced to

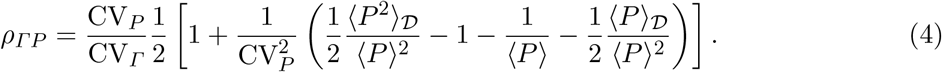

To relate this result to the mRNA–protein correlation, we must further specify what sets the total protein production rate, *Γ*.

### 3.3 mRNA-protein correlation for same-gene pairs

The protein production rate may depend on different factors, including mRNA abundance or ribosome availability [7, 29]. Before turning to a more detailed analysis, we consider here arguably the simplest translation model for the same-gene pair. Within this model, the protein production rate *Γ* is proportional to the number of mRNAs, *M*, and the proportionality coefficient is constant throughout the cell cycle (but may depend on the growth condition) [2, 5, 25]. Other translation models are discussed in Appendix Sec. C.

Combining this *Γ ∝ M* assumption with Eq. (4) yields

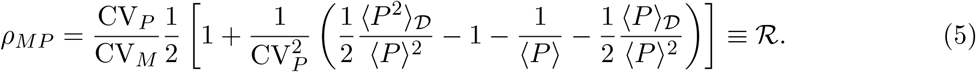

We test this mRNA–protein correlation using the Gillespie simulation [28] across several gene circuits (Fig. 2(d)). In these simulations, cells grow and divide deterministically, with stochastic binomial partitioning of cellular components at division. As shown in Fig. 2(e), Eq. (5) agrees with the stochastic simulations for all scenarios considered. The last term, 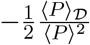, in Eq. (5) accounts for binomial partitioning at cell division; omitting this term reduces the equation to the corresponding result for deterministic equal partitioning. At large copy numbers, the two predictions are indistinguishable (Appendix Fig. A.1 (b) for ⟨*P* ⟩ ≃ 100), indicating that the effect of binomial partitioning is negligible in this limit. At low copy numbers, however, the deterministic-equal-partitioning approximation deviates from the binomial-partitioning result as expected (Appendix Fig. A.1 (a) for ⟨*P* ⟩ ≃ 10).

Under the same assumption *Γ ∝ M* and in the limit *λ* → 0, with a finite mean protein lifetime 1*/γ*_*p*_, Eq. (3) reduces to Eq. (1), thereby recovering the expression previously derived for mRNA– protein correlations in constant-volume, non-dividing cells [8]. In contrast to Eq. (5), Eq. (1) fails to predict the mRNA–protein copy numbers and concentration correlations in growing and dividing cells (see Fig. 2(b)).

### 3.4 mRNA-protein correlation for different-gene pairs

For different-gene pairs, if mRNA and protein neither regulate one another (not even indirectly) nor share upstream regulation, and if cell volume is their sole common source of correlation, then the correlation between *M* and *P* is given by

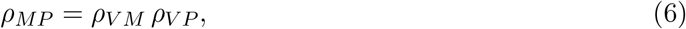

provided that both variables depend linearly on cell volume, *V*, with residual fluctuations that are uncorrelated with one another and with *V* [30]. Here, *ρ*_*V M*_ and *ρ*_*V P*_ denote the mRNA–volume and protein–volume correlations, respectively. We compare Eq. (6) for different-gene pairs against stochastic simulations and find good agreement (see Appendix Fig. E.1).

### 3.5 Comparison with experimental data

We test the predicted mRNA–protein correlations against the single-cell *E. coli* measurements [11] introduced in Sec. (3.1). For same-gene pairs, the measured mRNA–protein correlations, both with and without feedback, are consistent with the invariant prediction of Eq. (5) (blue circles in Fig. 3). None of these points deviates significantly from the identity line; after Bonferroni correction, all *p*-values are equal to 1. For different-gene pairs, the measured correlations are likewise consistent with the volume-mediated prediction of Eq. (6) (red squares in Fig. 3(b)). Again, none of these points deviates significantly from the identity line, with all Bonferroni-corrected *p*-values exceeding 0.05.

**Figure 3:**
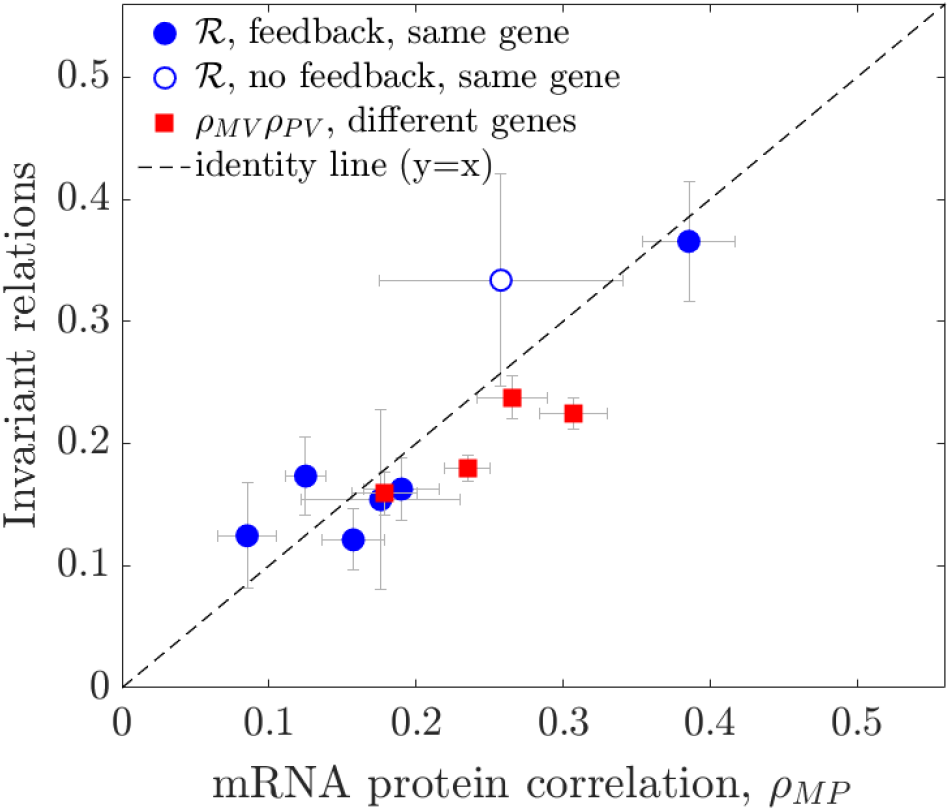
Experimental validation of mRNA–protein correlations predicted by Eq. (5) and (6). For the same experiments introduced in Sec. 3.1, mRNA–protein copy number correlations for different-gene pairs (red squares) are well described by Eq. (6), whereas Eq. (5) accurately reproduces the measured correlations for same-gene mRNA–protein pairs (blue circles). Error bars indicate bootstrap standard deviations obtained by resampling individual cells. The dashed line indicates *y* = *x* (perfect agreement between theory and experiment). Estimating R from Eq. (5) requires ⟨*P* ⟩_𝒟_ and ⟨*P* ^2^⟩_𝒟_, which cannot be obtained directly from the experimental data. We infer them from the protein time series using the Bayesian procedure described in Appendix Sec. B, and estimate the remaining mRNA/protein moments directly from the data.

To further examine the key assumption behind Eq. (6)—namely, that cell volume is the only confounder for different-gene pairs—we compute correlations between mRNA and protein *concentrations* (Appendix Fig. D.1). As expected, for same-gene pairs, the concentration correlations remain positive; whereas for different-gene pairs, the concentration correlations drop markedly. For two of the conditions, their value are statistically indistinguishable from zero (*p >* 0.1). For the other two, it is significantly positive (*p <* 10^−6^), indicating that cell volume is not the only confounding factor, at least for these two conditions (see Appendix Fig. D.1).

So far, we have adopted the simplifying assumption that the protein production rate is proportional to mRNA copy number, *Γ ∝ M*, leading to Eq. (5). More generally, translation may be limited by other factors, such as ribosome availability and mRNA fraction in the total mRNA pool, while transcription may be limited by gene copy number or RNAP copy number and gene fraction, giving rise to distinct gene-expression regimes [7, 29], as discussed in detail in Appendix Sec. C. We find that under fast-growth conditions, Eq. (5) holds in all regimes. Under slow-growth conditions, however, it is expected to fail in one regime, where transcription is limited solely by gene copy number, and translation is limited by both ribosome availability and mRNA fraction. However, the experimental data in Ref. [11] were collected in fast growth conditions, where the approximation *Γ ∝ M*, hence, Eq. (5) is also expected to hold. Discriminating among translation-limited regimes using single-cell mRNA-protein data, therefore, requires measurements at slower growth conditions.

### 3.6 Measurement noise

In addition to intrinsic noise, measurement noise can induce an additional source of variability in mRNA–protein correlations that was not included in the preceding analysis. Incorporating measurement noise yields a modified invariant relation for the correlation between the measured mRNA abundance, 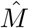, and the measured protein abundance, 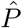, namely 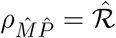 (see Eq. (A.33) in Appendix Sec. A.3).

Importantly, although mRNA measurement noise reduces the observed mRNA–protein correlation, it does not alter the invariant relation in Eq. (5), because Eq. (A.33) reduces to Eq. (5) in the absence of protein measurement noise (see Appendix Sec. A.3). As shown in Appendix Fig. A.2(a), multiplicative measurement noise in protein copy number reduces the observed correlation 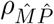 relative to the noiseless prediction of Eq. (5), thereby generating a systematic deviation from that relation. Nevertheless, Eq. (A.33) accurately predicts the correlation over a broad range of protein measurement noise levels, as illustrated in Appendix Fig. A.2(b).

### 3.7 Calibration of protein copy number

Absolute quantification of protein copy number in single cells remains technically challenging and depends critically on accurate calibration [31]. Importantly, Eq. (5) is not invariant under a rescaling of *P*. If the true protein copy number *P* is related to the measured, uncalibrated signal 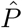 by 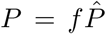, where *f* is a constant calibration factor, then this rescaling leaves the mRNA– protein correlation coefficient *ρ*_*MP*_ unchanged. By contrast, the theoretical prediction ℛ in Eq. (5) depends explicitly on *f* through terms proportional to 1*/*⟨*P* ⟩ and ⟨*P* ⟩_𝒟_*/*⟨*P* ⟩2. Consequently, an incorrect calibration factor introduces a systematic discrepancy between the predicted and observed correlations. This sensitivity enables Eq. (5) to be used to infer the protein calibration factor.

To quantify the agreement between theory and data (simulation and experiment), we compute, for each value of *f*, the root-mean-square (RMS) error, 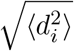, where *d*_*i*_ = ℛ− *ρ*_*MP*_ is the residual for data point *i*, defined as the difference between the theoretical prediction and the measured correlation. The calibration factor is then inferred as the value *f*_min_ that minimizes the RMS error. To estimate the uncertainty in *f*_min_, we performed bootstrap resampling of the individual sample points. We used the resulting distribution of *f*_min_ to determine its 95% bootstrap confidence interval, shown as an error bar in Fig. A.3.

Applying this procedure to simulated data yields *f*_min_ = 1.07, with a 95% confidence interval of 0.92–1.30, as expected for correctly calibrated protein copy numbers (Appendix Fig. A.3(a)). However, for *E. coli* data [11] we obtain *f*_min_ = 0.52, with a substantially broader 95% confidence interval of 0.36–1.07 (Appendix Fig. A.3(b)). This broader interval likely reflects both the limited number of data points (*n* = 7) and their relatively large uncertainty, suggesting that additional measurements and improved precision should substantially sharpen the estimate of *f*. Moreover, the value of *f* that we find here falls within the estimated error of the calibration procedure in [11].

### 3.8 Invariant relation in a mother-machine setup

So far, we have considered a population in which each division produces two daughter cells that both remain in the population. We now contrast this with a *lineage* setting, motivated by mothermachine experiments: a single “mother” cell is trapped at the closed end of a growth channel, while its newborn progeny are displaced by growth and washed out, enabling long-term tracking of a single lineage [32]. For this lineage ensemble, we obtain the following invariant relation (see Appendix Sec. F):

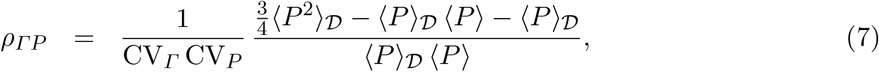

which differs from the invariant relation in Eq. (4), derived for the scenario where both daughter cells remain after division in the population or other systems with random cell dilution. This difference indicates that the invariant relation is ensemble-dependent, reflecting the distinct age structure and sampling statistics that affect mRNA–protein correlations.

To connect Eq. (7) to mRNA–protein correlations, we specify the total protein production rate *Γ* for the three gene-expression regimes discussed in Appendix Sec. C. The qualitative picture remains unchanged. Under fast-growth conditions, all three regimes become effectively indistinguishable and collapse onto the same invariant relation (Appendix Fig. F.1(a)). For slow-growth conditions, in the regime where transcription is limited by gene copy number only, and translation is limited by both ribosomes and mRNA fraction, a distinct relation is valid (see Appendix Sec. F).

## 4 Discussion

Quantitative models of complex biological systems often involve many interacting components with poorly constrained parameters [33]. This “curse of dimensionality” can increase the risk of overfitting empirical data [34]. An alternative strategy is to focus on a subsystem of interest, introducing simplifying assumptions about the remaining components and testing the robustness of the resulting conclusions to these assumptions [33]. Only in rare cases can exact invariant relations be derived—relations that depend solely on the subsystem of interest and remain insensitive to the details of the rest of the system [8]. An analogous situation arises in statistical mechanics, where a small number of universal relations, such as the Jarzynski equality [35], apply to broad classes of nonequilibrium systems despite widely differing microscopic dynamics.

In this work, we extended the framework of Ref. [8] to growing and dividing cells and derived invariant relations connecting the covariance between an mRNA and its protein product to their first and second moments. A key result is that these relations depend only on the details of translation and remain insensitive to the potentially complex upstream transcriptional dynamics.

At the same time, we showed that cellular growth and division qualitatively modify the form of these invariants. In particular, we obtained distinct results for *(i)* populations of cells with constant size, which recover the results of Ref. [8]; *(ii)* growing and dividing cells subject either to no removal or to stochastic removal of cells at division; and *(iii)* growing and dividing cells in which one daughter cell is always removed, as in mother-machine experiments [32].

Throughout, we considered a translation model in which the protein production rate scales linearly with mRNA abundance across all regimes examined in Sec. C. Although this assumption is widely used [2, 5, 7, 25], it is not universally valid; for example, a finite ribosome pool may produce a sublinear, Michaelis–Menten-like dependence of protein production on mRNA levels [36]. By analyzing single-gene measurements using the invariant relation derived here, we find that the linear approximation is consistent with the available empirical data, while noting that it need not hold for other genes, organisms, or growth conditions. Finally, our analysis focused on relations between the moments of absolute mRNA and protein copy numbers. In many contexts, however, concentrations are the more relevant observables. Unlike systems of constant volume, for growing and dividing cells, the statistics of concentrations can differ substantially from those of absolute copy numbers. Likewise, conditioning on cell volume may strongly reduce, or even eliminate, apparent correlations between mRNA and protein abundances, as in Ref. [37]. Extending the present framework to more general settings, including concentration-based statistics and volume-conditioned correlations, is therefore a natural direction for future work.

Recent advances in fluorescent protein labeling [38] and mRNA quantification technique [39, 40, 41], combined with the invariant relations derived here, may provide a useful framework for addressing quantitative aspects of the central dogma. Because these relations hold under broad conditions and depend only on the effective translation dynamics, they can serve as consistency checks for synthetic genetic circuits and experimental measurement pipelines. In particular, deviations from the predicted invariant relations may indicate translation dynamics in which protein production is not proportional to mRNA copy number or non-binary fission, neither of which is incorporated into the present analysis. Moreover, as demonstrated in Sections 3.6 and 3.7, the predicted invariant relation can be leveraged to estimate measurement noise and to calibrate experimental readouts, thereby enabling the conversion of fluorescence intensity into absolute protein copy numbers. More generally, our results provide a principled approach for probing translation dynamics within arbitrary transcriptional circuits without requiring detailed knowledge of the upstream regulatory architecture. In this sense, the framework developed here may be viewed not only as a tool for studying mRNA–protein coupling, but also as a more general strategy for isolating and characterizing a reaction of interest embedded in a complex and only partially observed interaction network in growing and dividing cells.

## Acknowledgments

K.B., M.S., and A.A. were supported by the European Union (ERC, BIGR, 101125981) and the Israeli Science Foundation (146873). I.G. is supported by the National Institutes of Health grant R35 GM140709, the National Science Foundation grant 2243257 (NSF Science and Technology Center for Quantitative Cell Biology), and the Alfred P. Sloan Foundation grant G-2023-19649.

## A Calculation of invariant relations for a population of growing and dividing cells

Here we derive Eqs. (3) and (F.15) building on the formalism developed in [8]. As mentioned in the main text, we consider a population of growing and dividing cells into two daughter cells, forming a tree with or without random cell dilution. Within the cells, we consider the expression of a gene embedded in an arbitrary, unspecified network that affects transcription dynamics, while we specify the translation process [8]. We make no assumptions about the growth process or the division-control mechanism. We start from the Markovian master equation for the probability density of the cell state described by the vector of numbers of all its components

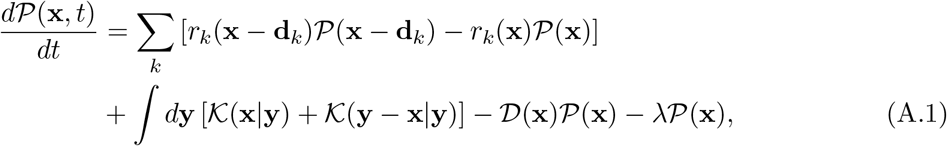

where we assume binomial partitioning of cell components

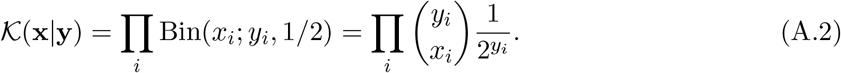

Previously mentioned in the main text, *r*_*k*_(**x** − **d**_*k*_) 𝒫 (**x** − **d**_*k*_) is the probability flux from state **x** − **d**_*k*_ to state **x** and 𝒟(**x**) is the division rate at state **x**. For the deterministic scenario, 𝒟 (**x**) will be a delta function. The sum runs over all possible transitions *k* and each transition is represented by an integer vector **d**_*k*_. To keep the normalization ∫𝒫 (**x**, *t*)*d***x** = 1 we remove random cells from the population with the growth rate *λ*, resulting in the −*λ*𝒫 (**x**) term.

The population averages ⟨*x*_*i*_⟩ (*t*) = 𝒫 (**x**, *t*)*x*_*i*_*d***x** follow

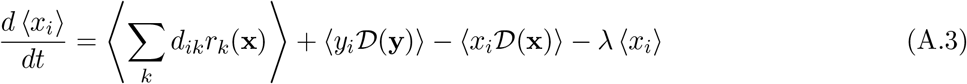

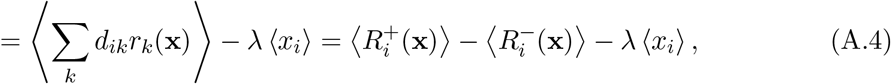

where we used 𝒦(**x**|**y**) = 𝒦(**y** − **x**|**y**), ⟨*x*_*i*_ 𝒦(**x**|**y**)⟩ = *y*_*i*_*/*2 and

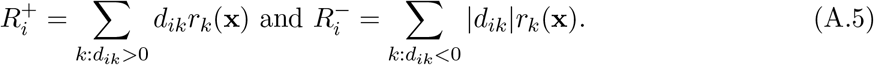

Here *d*_*ik*_ is the *i*’th element of the **d**_*k*_ vector. At the stationarity, 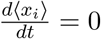 we have a bias towards the positive rate to balance the cell divisions:

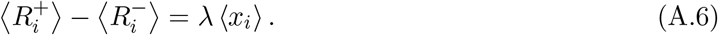

To calculate the second moments we use

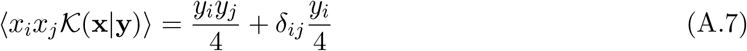

and get

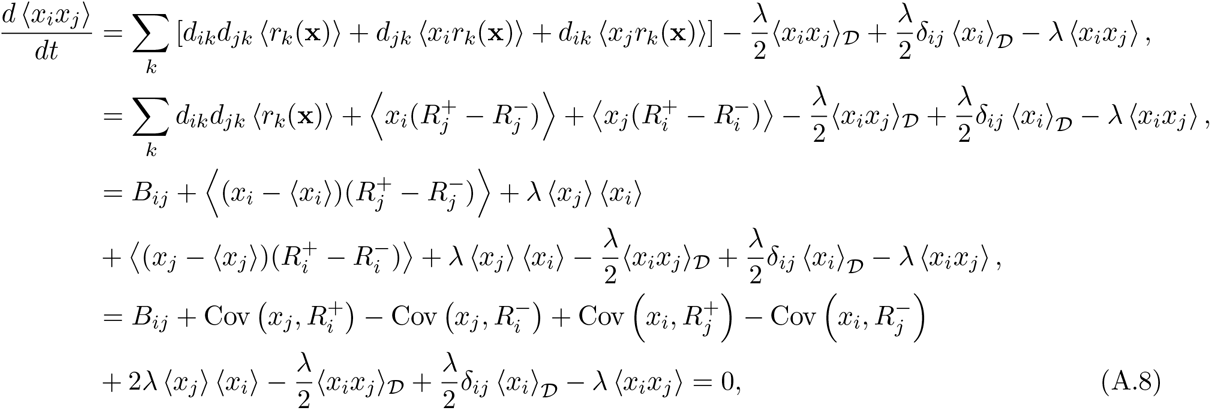

where

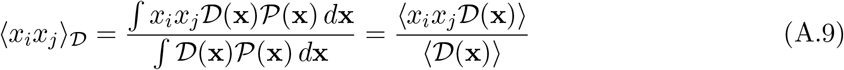

denotes the average value of *x*_*i*_*x*_*j*_ at division. The normalization 𝒟(**x**)𝒫(**x**) *d***x** = ⟨𝒟(**x**)⟩ = *λ* corresponds to the population growth rate. With this definition, the contribution 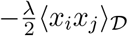 in Eq. (A.8) arises from the term ⟨2*x*_*i*_*x*_*j*_𝒟(2**x**)⟩ − ⟨*x*_*i*_*x*_*j*_𝒟(**x**)⟩.

The diffusion matrix can be written as

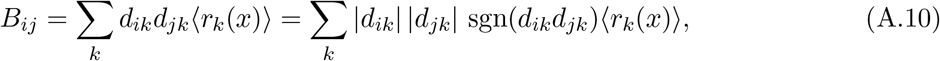

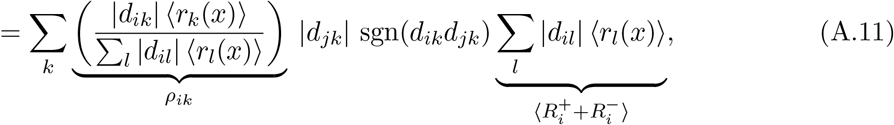

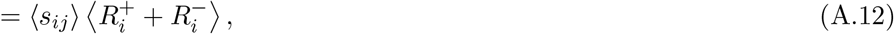

by defining

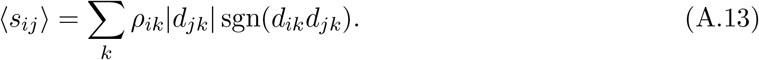

In sum, we get

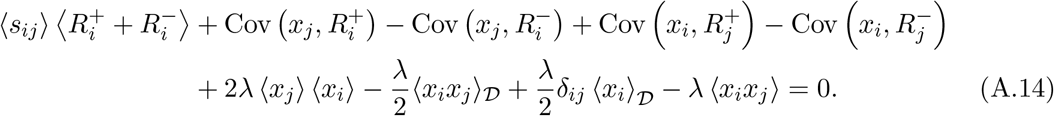

Using

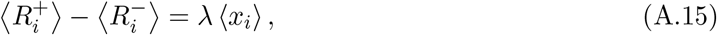

one gets

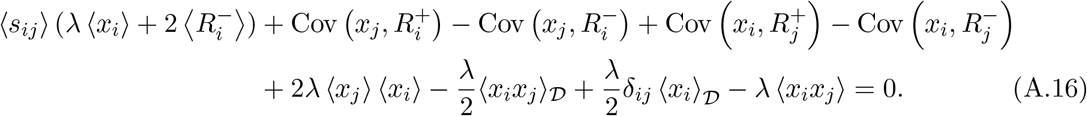

Assuming that all processes except division change the number of all elements by ± 1 we get ⟨*s*_*ij*_⟩ = 1. Denoting average degradation time of *x*_*i*_ as *τ*_*i*_ and using Little’s law [20] we get 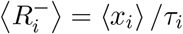.

This results in

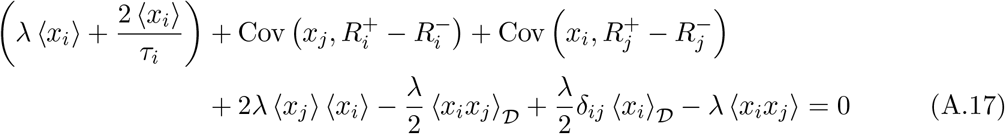

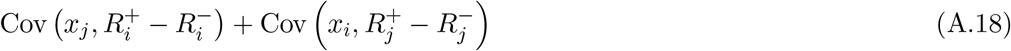

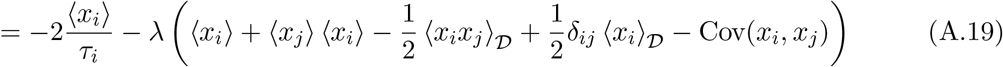

For *i* = *j* we get

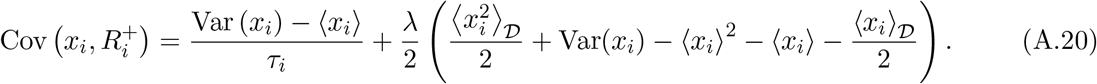

Consider now a more specific situation where index 1 denotes mRNA, *x*_1_ = *M*, and index 2 denotes protein translated from this mRNA, *x*_2_ = *P*. Assuming a constant protein degradation rate *γ*_*p*_ and some *M* dependent protein production rate 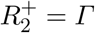 we have

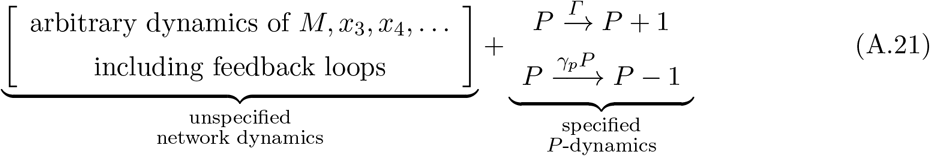

From Eq. (A.15), substituting *i* = 2 and *x*_2_ = *P*, we get

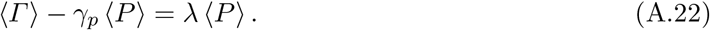

Using this, we get the correlation between the protein copy number *P* and the total protein production rate *Γ* as follows

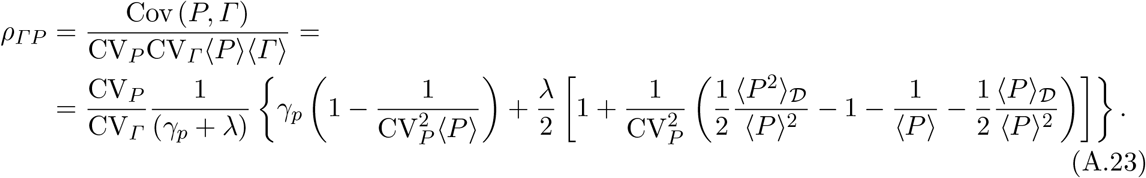

For *γ*_*p*_ ≪ *λ*, Eq. (A.23) simplified to

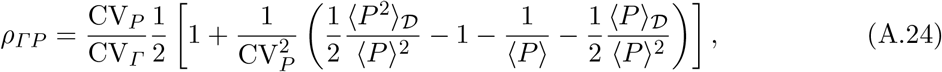

which is the same as Eq. (4) in the main text. Eq. (A.24) is invariant in the sense that it does not depend on the transcription, translation, degradation, genome replication, or division control. Next, we will discuss different forms of invariant relation in Eq. (A.24) for different scenarios of protein production rate *Γ*.

### A.1 Invariant relation for mRNA copy number limited translation

If the protein production rate *Γ* is proportional to the number of mRNAs, *M*, one gets the following dynamics

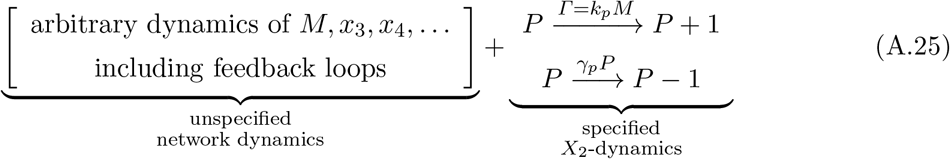

Using Eq. (A.24) with *Γ ∝ M* we obtain

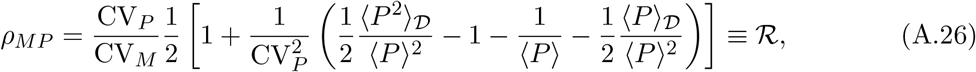

which is the same mRNA–protein correlation in the main text in Eq. (5).

**Figure A.1:**
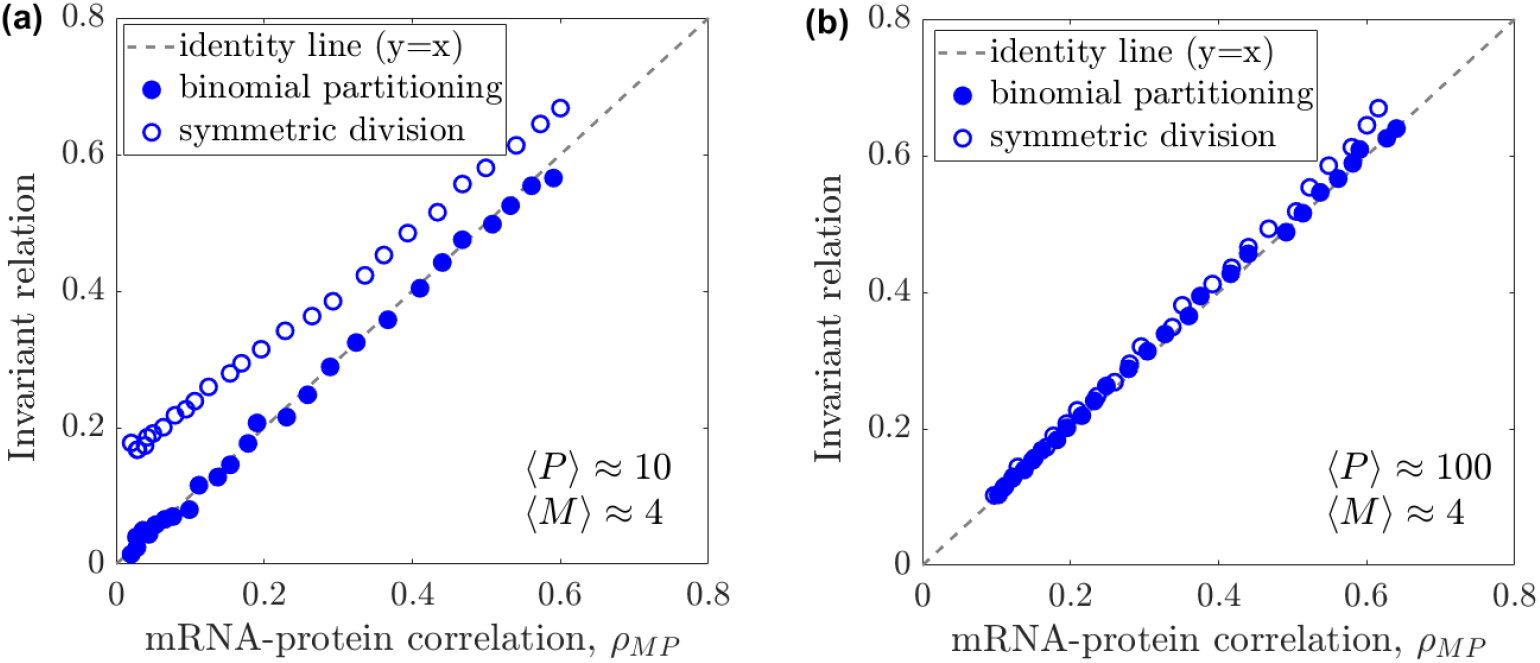
mRNA-protein correlation for symmetric and binomial partitioning. The mRNA–protein correlation, *ρ*_*MP*_, obtained from stochastic simulations with binomial partitioning at cell division, is plotted against the corresponding invariant prediction. Filled symbols show the prediction of Eq. (A.26) including the binomial-partitioning correction, whereas open symbols show the corresponding prediction obtained by omitting the last term, 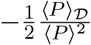 in Eq. (A.26), i.e., the approximation for deterministic equal partitioning at division. **(a)** In the low-copy-number regime, the full invariant prediction, including binomial partitioning, remains in good agreement with the simulation results, whereas the deterministic-equal-partitioning approximation shows a clear deviation. **(b)** In the high-copy-number regime, the two predictions become nearly indistinguishable, showing that the correction due to binomial partitioning becomes negligible at large copy numbers. The symbols are obtained from stochastic simulation [28] over 10^3^ cells.

Stochastic binomial partitioning at cell division reduces correlations between cellular components relative to deterministic symmetric division. This reduction is set by the variance of the binomial distribution and is captured by the 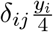 term in Eq. (A.7). In turn, this contribution generates the 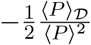 terms in Eqs. (A.23,A.24,A.26). Omitting this term, one obtains the expressions corresponding to deterministic symmetric cell division. As shown in Fig. A.1(b), the predicted correlations for binomial and symmetric partitioning become indistinguishable at large protein copy numbers, indicating that the correction due to binomial partitioning is negligible in this limit. At low protein copy numbers, however, the symmetric division deviates from the binomial-partitioning result as expected (Fig. A.1(a)).

### A.2 Invariant relation for ribosome and mRNA fraction limited translation

If the protein production rate is limited by the ribosome number and mRNA fraction, then 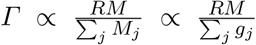, if the transcription rate is limited by the total gene dosage, such that 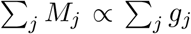. The denominator, Σ _*j*_ *g*_*j*_, scales with the total cellular DNA content *DNA*(*t*), which increases piecewise linearly and doubles once per cell cycle [42, 43]. Assuming that the number of ribosomes scales linearly with the volume *R ∝ V* (*t*), one gets the following dynamics

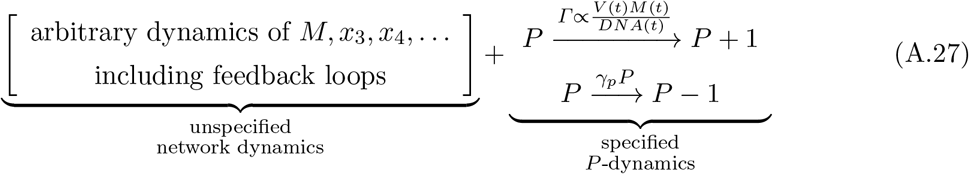

Using Eq. (A.24) with 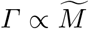, by defining 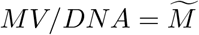, we obtain

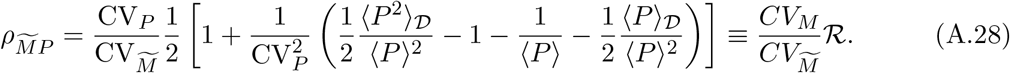

The function *DNA*(*t*) can be obtained from the Cooper-Helmstetter model. For *not* extremely slow-growing cells *DNA* ∼ *V*, such that 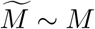, hence *ρ*_*MP*_ ≃ ℛ.

### A.3 Effects of measurement noise on mRNA–protein correlation

We consider multiplicative measurement noise affecting both mRNA and protein abundance, such that the measured mRNA and protein copy numbers are given by 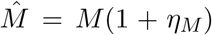 and 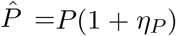, respectively. Here, *η*_*M*_ and *η*_*P*_ are independent Gaussian random variables with zero mean and variances 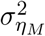 and 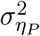. The observed mRNA–protein correlation is then

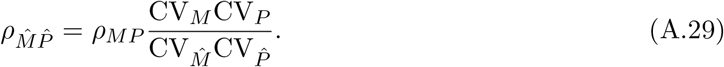

Substituting *ρ*_*MP*_ from Eq. (A.26) yields

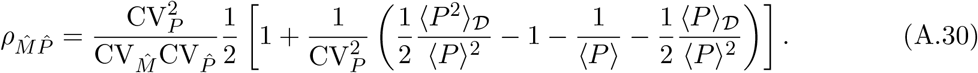

Using the relations for the second moments and coefficients of variation,

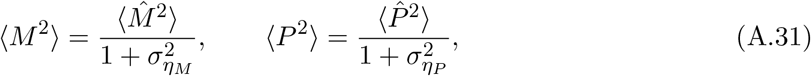

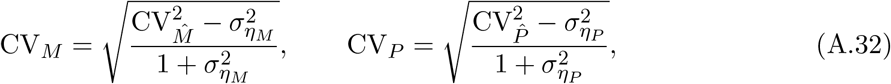

we obtain

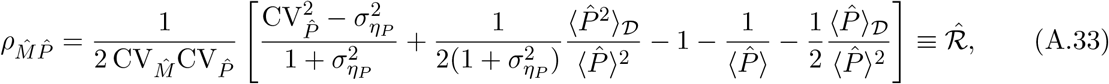

where we assume that the protein abundance measured at division is affected by multiplicative noise with the same variance 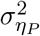. Notably, the invariant Eq. (A.33) reduces to Eq. (A.26) when protein measurement noise 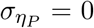, indicating that only protein measurement noise modifies the invariant relation. In other words, although measurement noise in mRNA abundance can influence the observed mRNA–protein correlation, such noise does not affect the invariant relation.

**Figure A.2:**
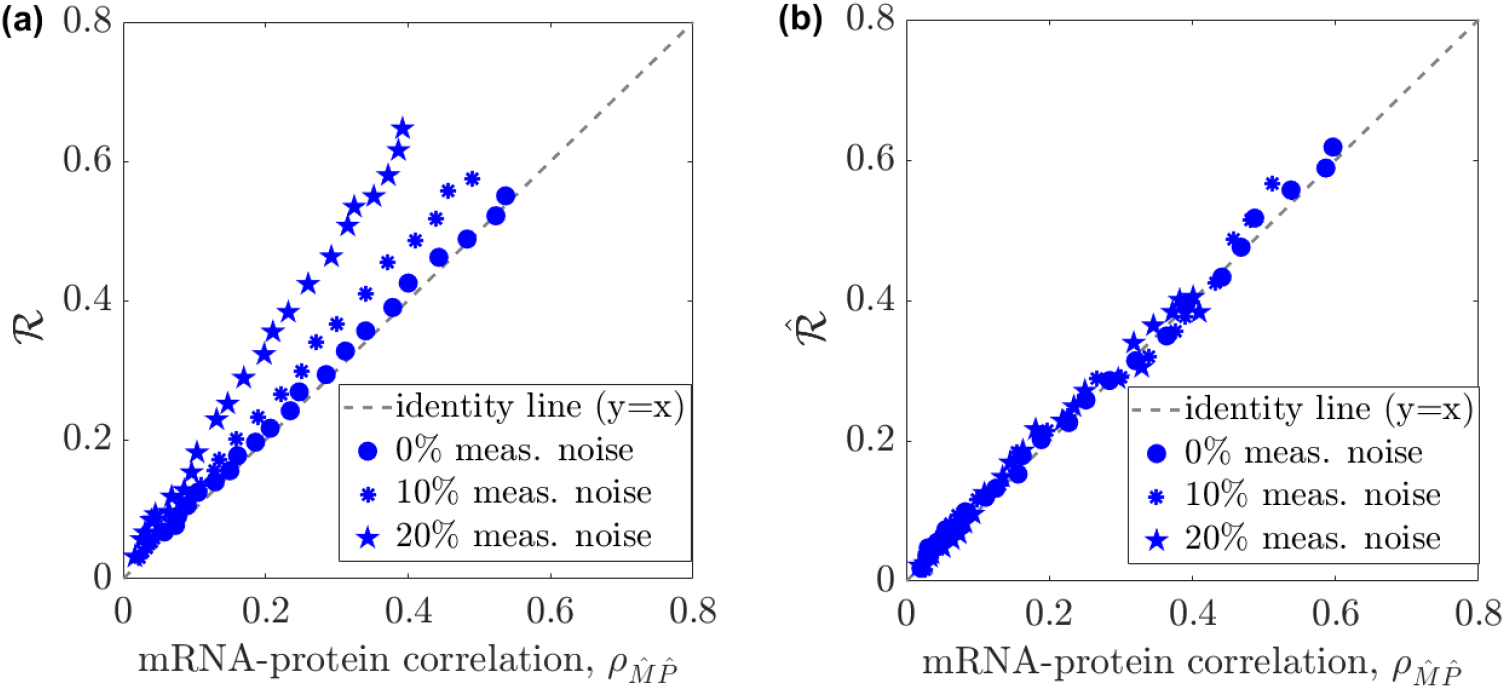
Measurement noise attenuates the observed mRNA–protein correlation. **(a)** In the presence of multiplicative measurement noise in protein copy numbers, the observed correlation 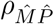 is reduced relative to the noiseless prediction of Eq. (A.26), resulting in a systematic deviation from the identity line. **(b)** The correlation in the presence of measurement noise is predicted by Eq. (A.33) and is in good agreement with simulation. Symbols denote different levels of measurement noise, and the dashed line indicates the identity line. The symbols are obtained from stochastic simulation [28] over 10^3^ cells.

### A.4 Calibration of protein copy number

**Figure A.3:**
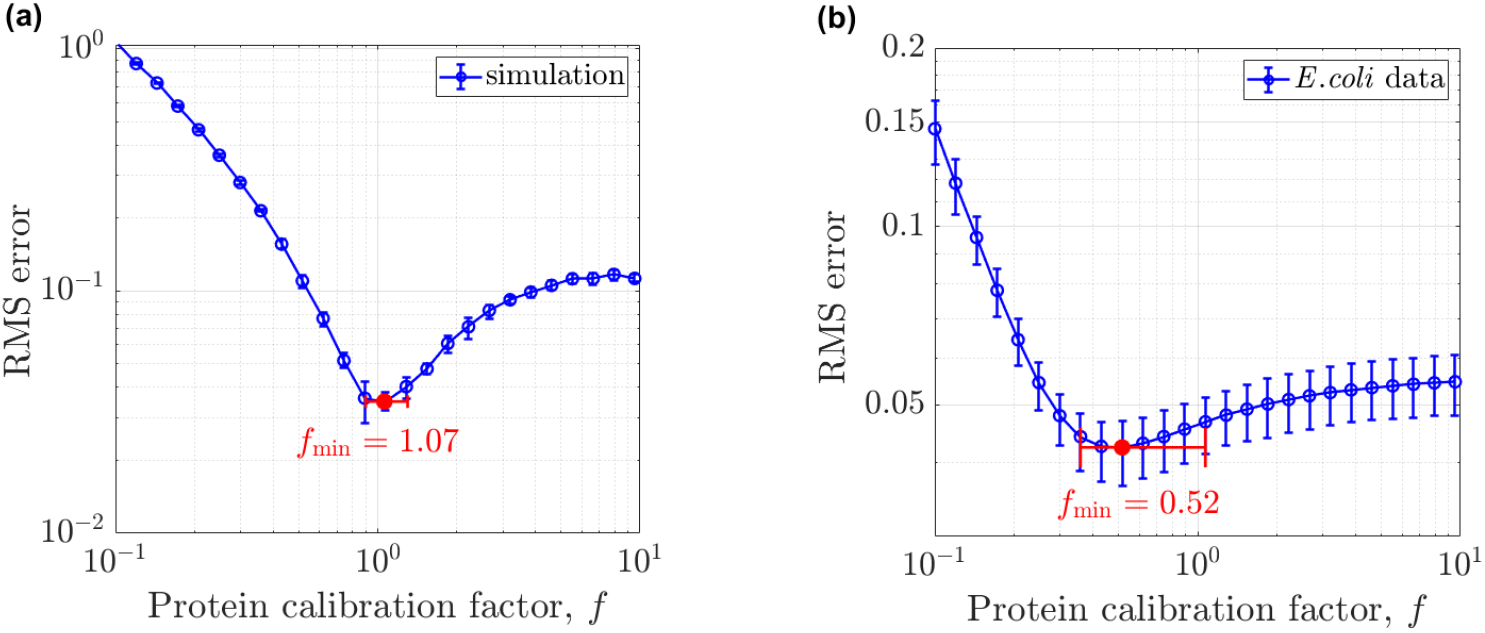
Inferring the protein copy-number calibration factor from Eq. (5). We consider that the true protein copy number *P* is related to the measured, uncalibrated signal 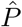 by 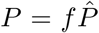, where *f* is a constant calibration factor. For each calibration factor *f*, we compute the root-mean-square (RMS) error for **(a)** simulated data and **(b)** *E. coli* data [11]. The RMS error is defined as 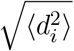, where *d*_*i*_ = ℛ−*ρ*_*MP*_ is the residual for data point *i. ρ*_*MP*_ is the measured mRNA– protein correlation, ℛ is the corresponding prediction from Eq. (5). The calibration factor is then inferred as the value *f*_min_ that minimizes the RMS error. **(a)** For simulated data, *f*_min_ = 1.07 with a narrow 95% confidence interval of 0.92–1.30. **(b)** For *E. coli* data, *f*_min_ = 0.52, with a substantially broader 95% confidence interval of 0.36–1.07. We performed bootstrap resampling of the individual sample points, and error bars are 95% bootstrap confidence interval.

## B Estimation of ⟨*P* ⟩_D_ and ⟨*P* ^2^⟩_D_

To validate the invariant relations in Eq. (5), we need to estimate *ρ*_*MP*_ and ℛ under different conditions. While *ρ*_*MP*_ can be obtained directly, estimating ℛ requires an accurate estimate of the first and second moments of protein number at division, respectively, denoted as ⟨*P* ⟩_*𝒟*_ and ⟨*P* ^2^⟩_*𝒟*_. However, directly measuring ⟨*P* ⟩_*𝒟*_ and⟨*P* ^2^⟩_*𝒟*_ is challenging from population-snapshot data. We therefore infer the protein-number statistics at division as follows.

We define ⟨*P*^*n*^⟩_*𝒟*_ as the *n*-th moment of the protein number at division:

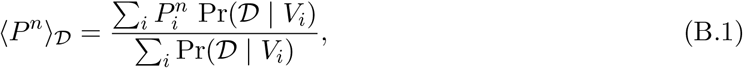

where Pr(𝒟 | *V*_*i*_) is the probability that a cell with volume *V*_*i*_ divides. Using Bayes’ theorem,

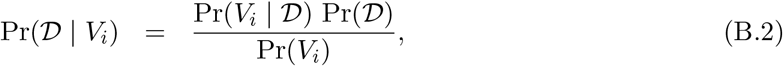

which yields

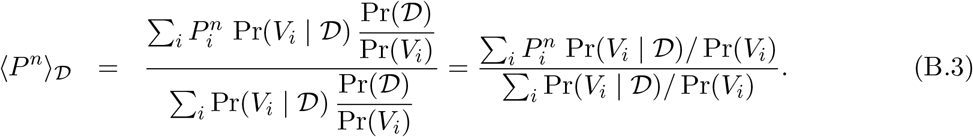

Unless stated otherwise, all probabilities in this section refer to population snapshots. The distribution of division volumes in the population, Pr(*V* | 𝒟), is given by

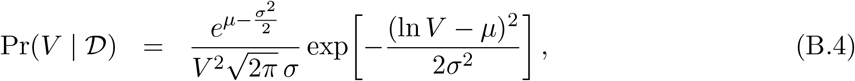

which follows from the relation

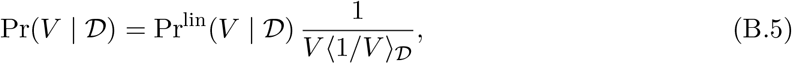

where Pr^lin^(*V* | 𝒟) is the lognormal distribution of division volume along a lineage [44]. Eq. (B.5) gives the relation between the division-volume distributions in the population and lineage ensembles, as reported in Refs. [19, 45].

For given parameters *µ* and *σ*, the snapshot distribution of cell volumes, Pr(*V*), is obtained from [19, 45]

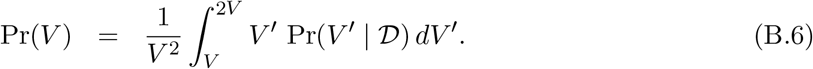

Note that in [19, 45] the distribution of division volumes is defined along lineages, whereas here we consistently express everything in terms of population-level probabilities. We estimate *µ* and *σ* by maximizing the log-likelihood of the observed snapshot volumes {*V*_*i*_},

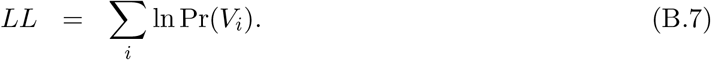

With the resulting estimates of *µ* and *σ*, we compute Pr(*V* | 𝒟) from Eq. (B.4), Pr(*V*) from Eq. (B.6), and subsequently ⟨*P* ⟩_*𝒟*_ and ⟨*P* ^2^⟩_*𝒟*_ from Eq. (B.3) with *n* = 1 and 2, respectively. We evaluate the performance of this inference procedure using simulated data as ground truth by comparing the inferred Pr(*V* | 𝒟) with the distribution measured directly from the simulations, and find good agreement (Fig. B.1(a)). We then extend the method to *E. coli* data to infer the population division-volume distribution, Pr(*V* | 𝒟), across 11 growth conditions and regulatory architectures, as shown in Fig. B.1(b–l).

**Figure B.1:**
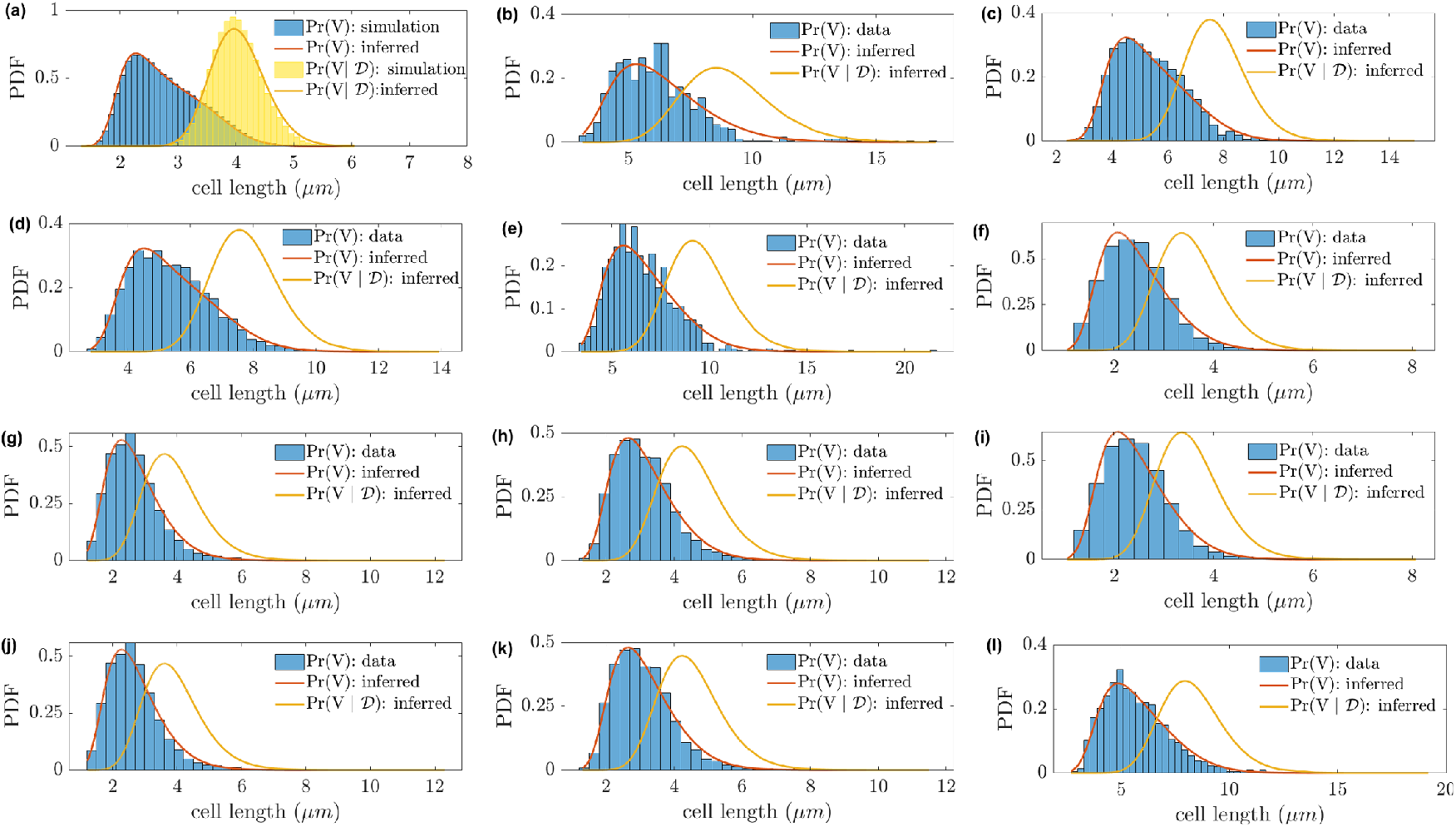
Inference of division-size distribution from snapshot data. **(a)** Validation using simulated data. Blue bars show the snapshot cell-size distribution, Pr(*V*), measured directly from the simulations, and the red curve shows the corresponding inferred distribution (Eq. (B.6)). Yellow bars show the division-size distribution in population, Pr(*V* | 𝒟), measured directly from the simulations, and the yellow curve shows the inferred distribution (Eq. (B.4)). Simulations were performed in a growing and dividing cell population, in which each cell elongated, divided into two daughter cells according to an adder size-control mechanism, and partitioned its contents binomially at division, with a constant division time of *T* = 50 min. **(b–l)** Application of the same inference procedure to *E. coli* data across 11 growth conditions and regulatory architectures. Blue histograms show the measured snapshot size distributions, the red curves show the inferred Pr(*V*), and the yellow curves show the inferred division-size distributions, Pr(*V* | 𝒟).

## C Invariant relations for different gene-expression regimes

Protein production is limited by mRNA and/or ribosome levels [7]. Their relative importance varies across conditions, giving rise to three physiological regimes [7, 29]. These gene-expression regimes are schematically illustrated in Fig. C.1(a). We adopt the nomenclature of Ref. [7] and analyze the invariant relation for each regime in turn.

### Regime III

First, we consider the regime in which transcription is limited only by gene dosage and translation is limited only by mRNA [29]. In this scenario, *Γ ∝ M*, and the invariant relation in Eq. (A.26) that we analyzed in Sec. A.1 is valid for this regime.

### Regimes I and II

In Regimes I and II, protein production is limited by both ribosomes, *R*, and mRNAs. For simplicity, similar to Refs. [7, 29], we assume that all mRNAs have comparable ribosome-binding properties and that ribosomes are allocated in proportion to each mRNA’s fraction of the total mRNA pool. Under these assumptions, the total protein production rate is given by

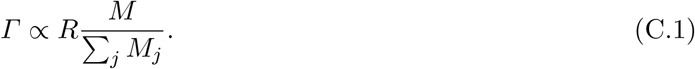

Here, *M*_*j*_ is the mRNA number of gene *j*, while *M* is the mRNA number of the gene of interest. Note that the total protein production rate in Eq. (C.1) differs from the simplified assumption *Γ ∝ M* used in Sec. A.1, because the prefactor *R/ Σ* _*j*_ *M*_*j*_ is not generally constant over the cell cycle. To specify this prefactor, we must characterize the temporal behavior of the total mRNA count, *Σ* _*j*_ *M*_*j*_. This behavior is set by the transcription-limiting factors that govern most genes (not necessarily the gene of interest). We therefore consider two regimes.

**Figure C.1:**
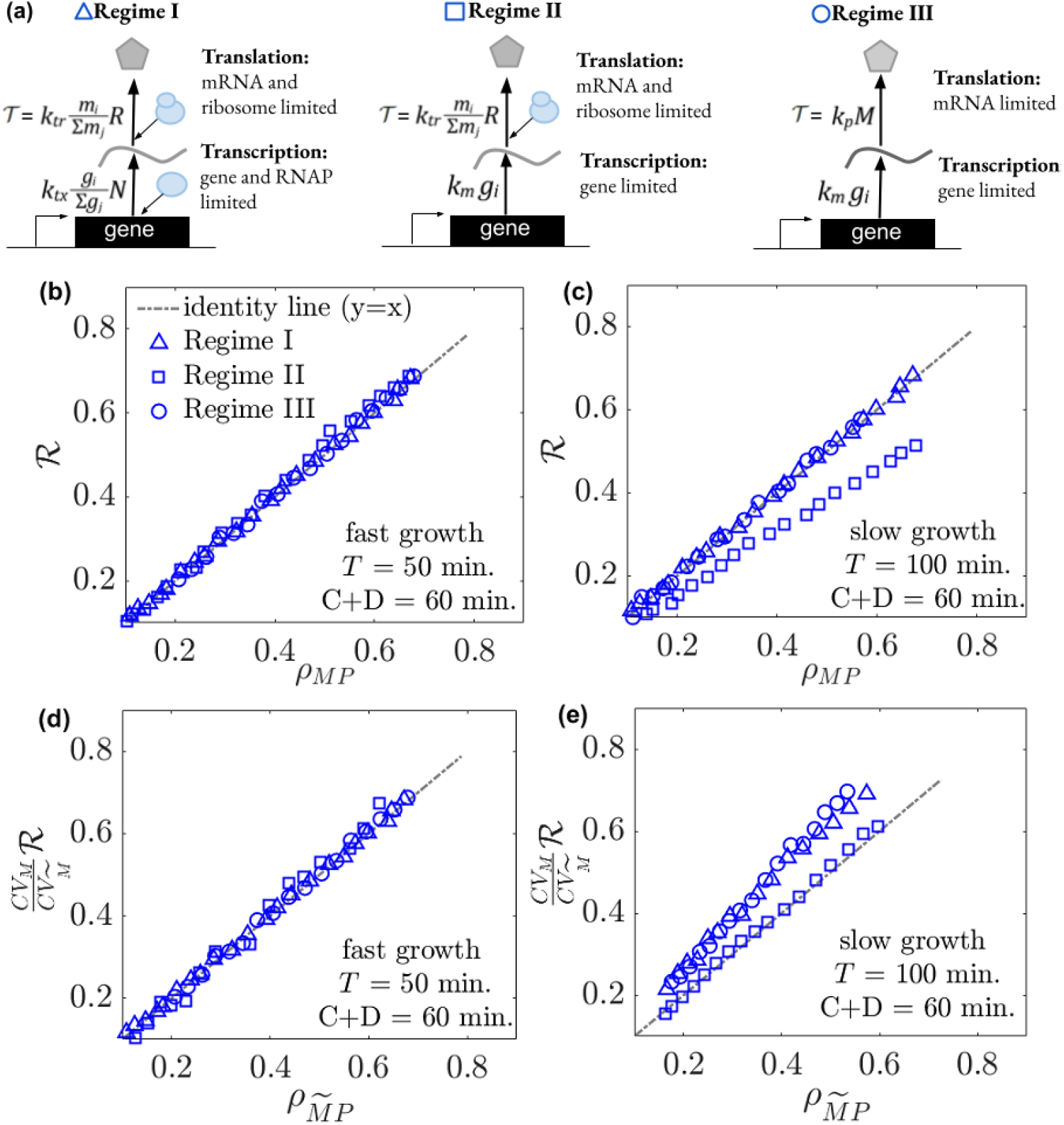
Using invariant relations to identify gene-expression regimes in slow growth. **(a)** Schematic of three different gene expression regimes adapted from [7], where protein and mRNA production rates are limited by different resources: transcription is limited by RNAPs or by the gene dosage, and translation is limited by ribosomes or by the mRNA. The definitions of various model parameters are provided in Appendix Table 1. The invariant Eq. (5) for all three regimes for **(b)** fast and for **(c)** slow growth conditions. In fast-growth (*C* + *D > T*), all regimes collapse to the same invariant relation, but Regime II deviates from invariant Eq. 5 for slow-growth condition (*C* + *D < T*). For Regime II, the mRNA–protein correlation measured from stochastic simulation follows the invariant Eq. (A.28) in both **(d)** fast and **(e)** slow growth conditions. However, other regimes deviate from the invariant Eq. (A.28) for slow growth conditions **(e)**. In all plots, symbols are from stochastic simulations [28] over 10^3^ cells. In all panels, the dashed line indicates the identity *y* = *x* for reference.

### Regime I

In this regime, the transcription across most genes is limited by the availability of both RNAP number (*N*) and gene-dosage (*g*_*i*_) fractions, resulting in *M*_*i*_ *∝ Ng*_*i*_*/ Σ* _*j*_*g*_*j*_. The total mRNA abundance, _*Σ i*_ *M*_*i*_ *∝ N ∝ V*, therefore scales proportionally with cell volume [46, 47]. Substituting this scaling into Eq. (C.1) yields *Γ ∝ M*, thereby recovering the same invariant relation (Eq. A.26) as in regime III and Sec. A.1.

### Regime II for fast-growing cells

If transcription for most genes is limited primarily by gene dosage, then the total mRNA abundance scales with the total gene copy number, Σ_*j*_ *M*_*j*_ *∝* Σ_*j*_ *g*_*j*_. Under this assumption, Eq. (C.1) is simplified to 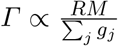. The denominator, Σ_*j*_ *g*_*j*_, scales with the total cellular DNA content, which increases piecewise linearly during chromosome replication and doubles once per cell cycle [42, 43]. For fast-growing cells, for which the doubling time *T* is shorter than *C* + *D* (where *C* is the chromosome replication period and *D* is the time between replication termination and cell division), the piecewise-linear increase in total gene copy number is, in principle, distinguishable from exponential growth given sufficiently precise measurements. Otherwise, when exponential and piecewise linear growths are experimentally indistinguishable, the ratio *R/ Σ*_*j*_ *g*_*j*_ is approximately constant throughout the cell cycle. In this case, we again obtain *Γ ∝ M*, as assumed in Sec. A.1, and recover the same invariant relation, Eq. (A.26), as in Regimes I and III.

### Regime II for slow-growing cells

In contrast, for slowly growing cells, where *T > C* + *D*, regime II deviates from Eq. A.26, because for a significant fraction of the cell cycle, total gene number *Σ*_*j*_ *g*_*j*_ does not grow, such that *R/ Σ*_*j*_*g*_*j*_ significantly deviates from a constant. In this case 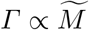, (where 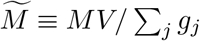), the same as in Sec. A.2, and recover, the invariant relation in Eq. (A.28).

Across all regimes considered, we validate the invariant relations using stochastic simulations [28] (see Fig. C.1). Under fast-growth conditions, when *T* ≲ *C* + *D*, all regimes conform to the invariant relation in Eq. (A.26) (Fig. C.1(b)). In contrast, under slow-growth conditions, *T > C* + *D*, Eq. (A.26) holds for Regimes I and III but not for Regime II (Fig. C.1(c)). In this slow-growth limit, Regime II instead satisfies the invariant relation in Eq. (A.28) (Fig. C.1(e)). These simulations therefore indicate that, at realistic copy numbers and the associated intrinsic noise levels, the regimes remain distinguishable under slow growth (Fig. C.1(c,e)) but are effectively indistinguishable under fast growth (Fig. C.1(b,d)).

## D mRNA–protein copy-number versus concentration correlations

**Figure D.1:**
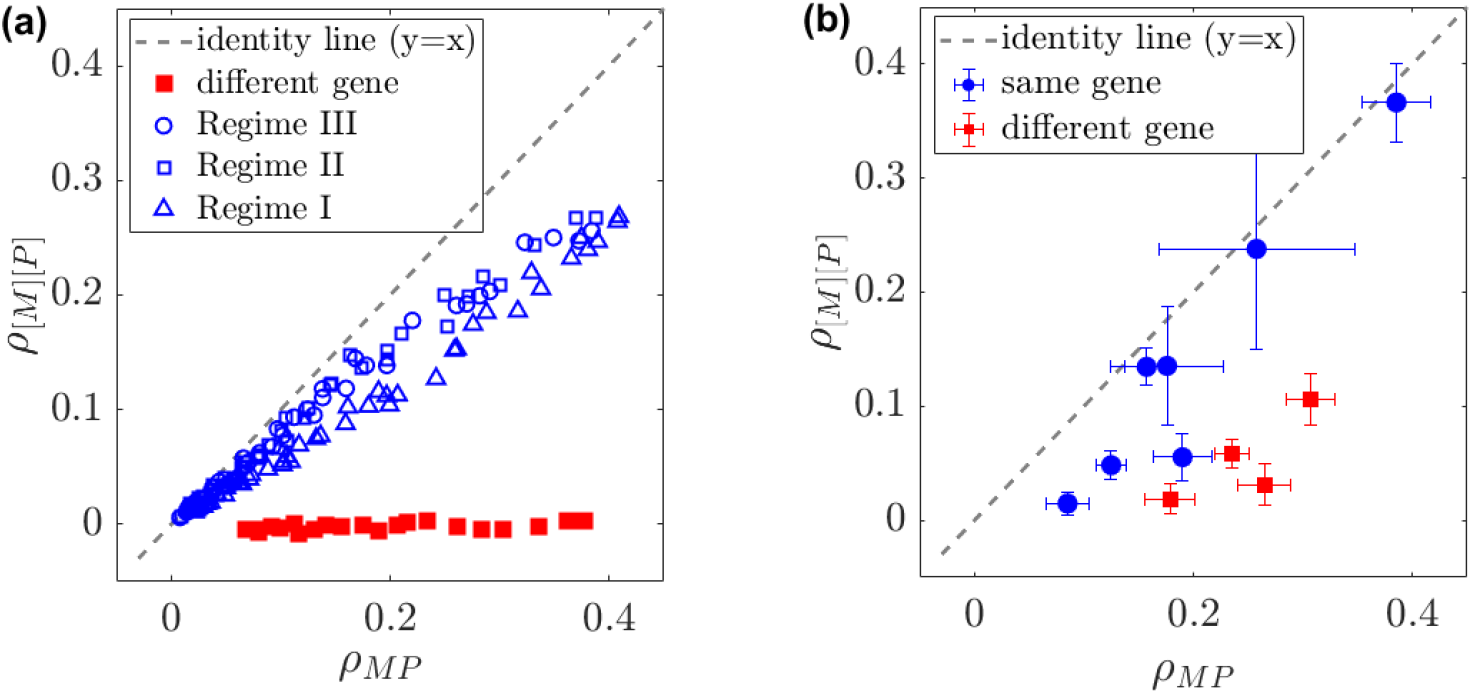
Correlation between mRNA and protein *concentrations* versus *copy numbers* in (a) simulations and (b) experimental data. **(a)** Simulated mRNA–protein concentrations correlation, *ρ*_[*M* ][*P* ]_, plotted against the corresponding copy-number correlation, *ρ*_*MP*_, for the same-gene (Regimes I–III; blue symbols) and different-gene pairs (red squares). The dashed line indicates *y* = *x*. For different-gene pairs, concentration correlations collapse toward zero because the copy-number correlation is mediated solely by cell-volume fluctuations. Same-gene pairs, however, retain a substantial positive concentration correlation, reflecting the direct dependence of protein production on mRNA. **(b)** The same analysis applied to *E. coli* experimental data [11] shows the same trend: concentration correlations remain positive for same-gene pairs, whereas they decrease markedly for different-gene pairs, while remaining above zero, indicating that cell volume is not the sole confounding factor.

## E mRNA-protein correlation for unrelated gene pairs

**Figure E.1:**
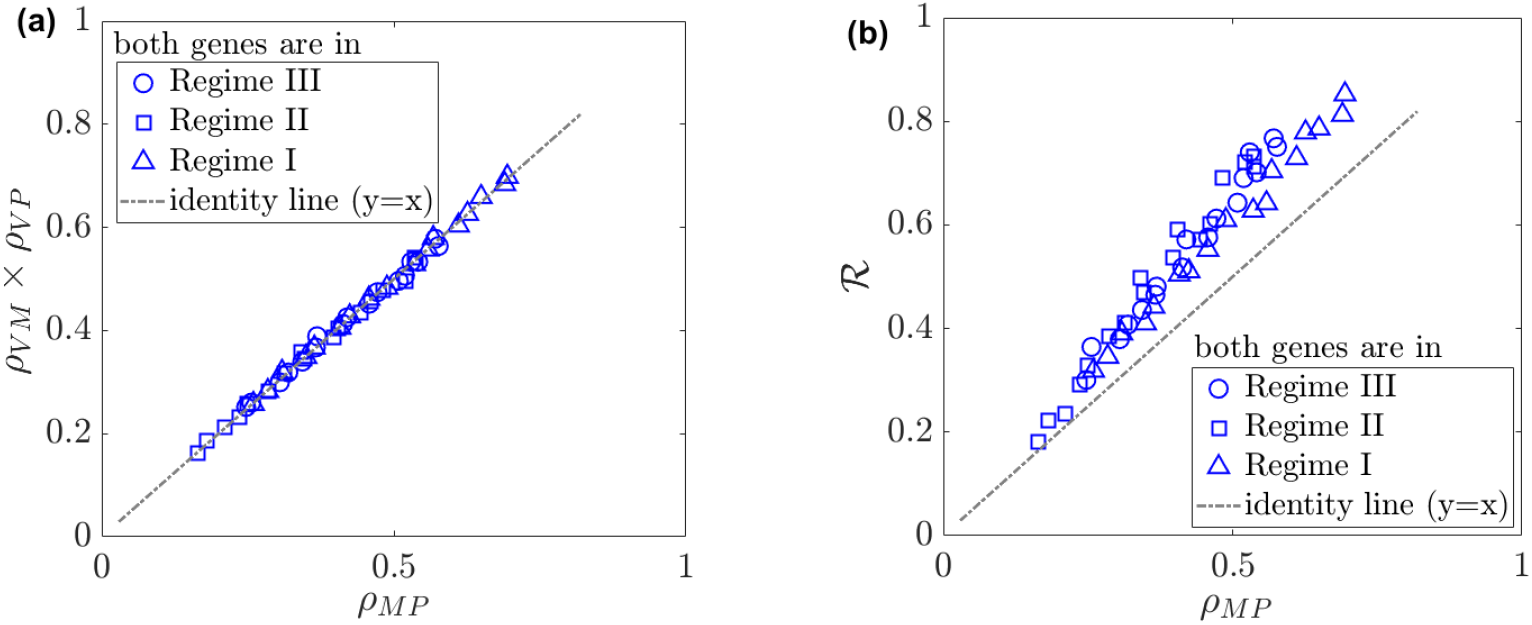
When mRNA and protein are from two uncorrelated genes, the mRNA–protein correlation reduces to the product of the mRNA–volume correlation (*ρ*_*V M*_) and the protein–volume correlation (*ρ*_*V P*_) because volume is the only confounding factor. This holds for all three gene expression regimes in Sec. C for *T* = 50 min. However, the invariant Eq. (5), *ρ*_*MP*_ = ℛ for the same gene pairs will overestimate the mRNA-protein correlation as expected. In all plots, symbols are from stochastic simulations [28] over 10^3^ cells. In both panels, the dashed line indicates the identity *y* = *x* for reference.

## F Calculation of invariant relations for a lineage of growing and dividing cells

In this scenario, relevant for a mother machine setup [32], a division event results in one daughter cell, the second one is removed from the population. This removal changes the 2 *d***y**𝒦(**x**|**y**) 𝒟(**y**)𝒫(**y**, *t*) term in master equation (2) (and Eq. (A.1) in Appendix) to *∫d***y**𝒦(**x**|**y**) 𝒟(**y**)𝒫(**y**, *t*) resulting in

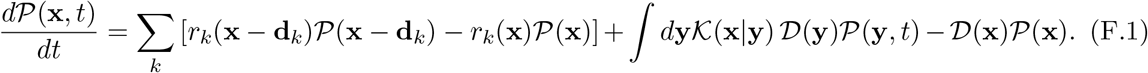

Using the same approach as in Section A, the averages follow

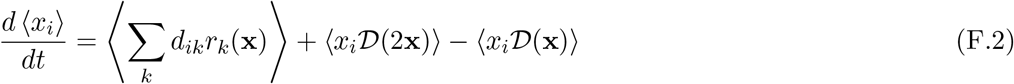

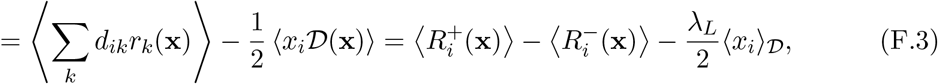

where ⟨*x*_*i*_⟩_*𝒟*_ is the average of *x*_*i*_ at the cell division and *λ*_*L*_ is the division rate averaged along the lineage

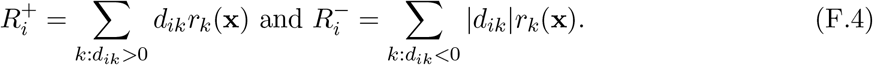

At the stationarity we have a bias towards the positive rate to balance the cell divisions:

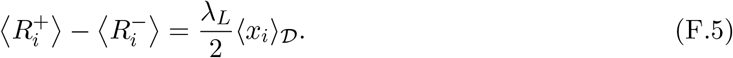

The second moments follow

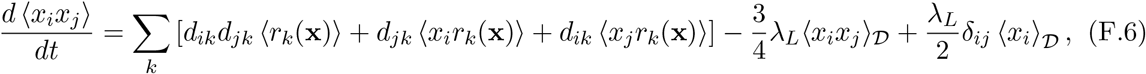

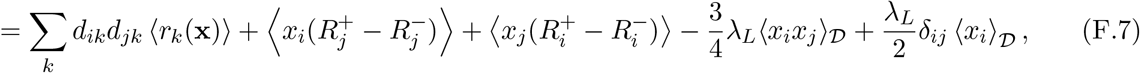

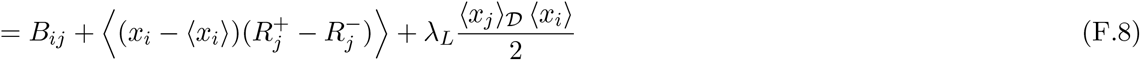

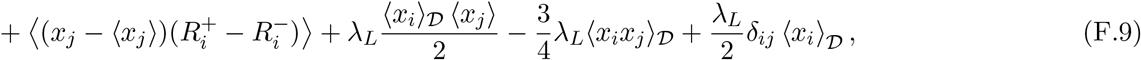

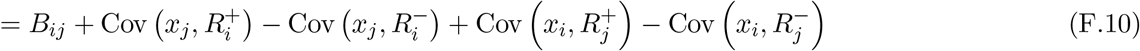

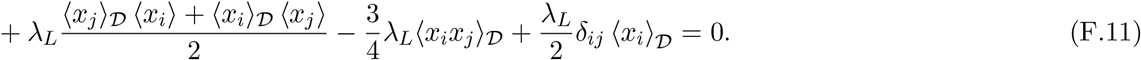

For *i = j* We get

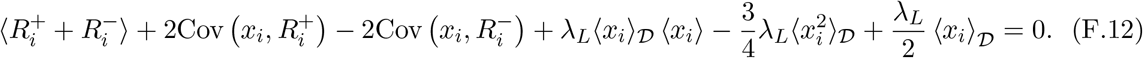

As before, for *i* = 2, 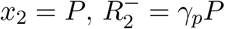 and 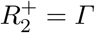 we get

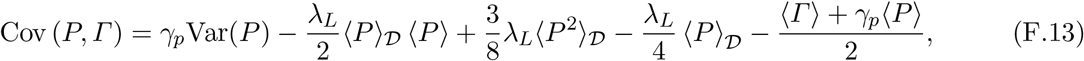

and using Eq. (F.5) 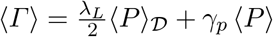

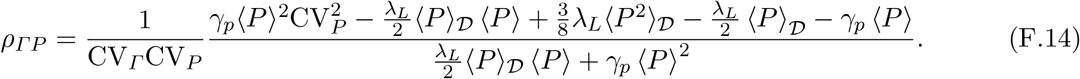

For *γ*_*p*_ ≪ *λ*_*L*_ this is simplified to

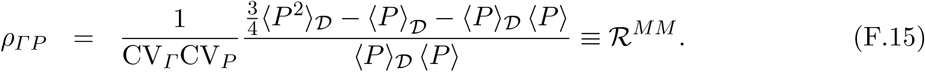

As before, for fast-growing cells, the invariant relation for all regimes

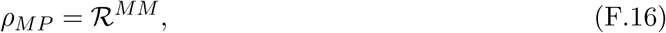

(see Fig. F.1(a)), and, otherwise, in regime II, the invariant relation,

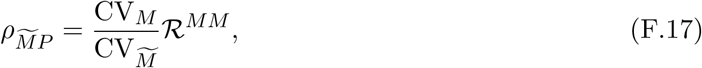

see Fig. F.1(b). It is important to note that the mRNA-protein correlation for the lineage in Eq. (F.15) follows a different invariant relation than that of the tree ensemble in Eq. (4) as shown in Fig. F.1(c). This distinction highlights that mRNA–protein correlations depend on the observational ensemble, and that predictions obtained for lineage measurements cannot, in general, be directly identified with those for tree-ensemble statistics.

**Figure F.1:**
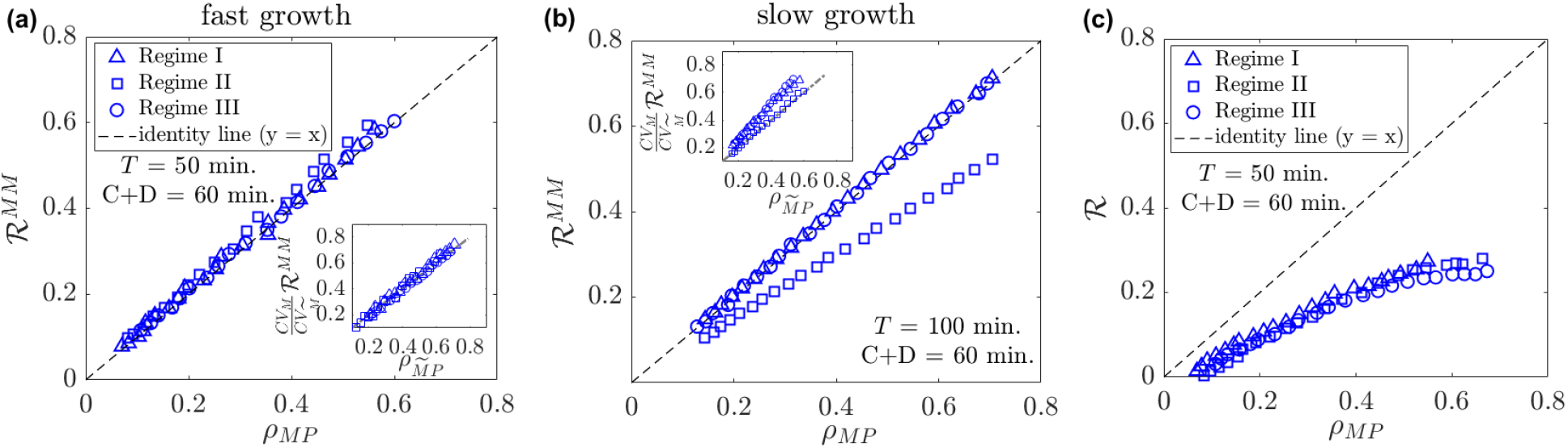
Invariant relations for a lineage across three different gene expression regimes. **(a)** For fast growth conditions, regimes are indistinguishable and follow the invariant relation in Eq. (F.16). Inset: all regimes will also follow the other invariant relation in Eq. (F.17) in the fast growth condition. **(b)** In the slow growth condition, Regime II will differ from invariant relation in Eq. (F.16) and follow the invariant relation in Eq. (F.17) (inset figure). However, Regimes I and III will differ from Eq. (F.17) in slow growth conditions (inset figure). **(c)** mRNA-protein correlation for cells within a lineage differs from the invariant relation for the tree ensemble in Eq. (5), indicating that the sampling method for cells matters for the mRNA-protein correlation. In all plots, symbols are from stochastic simulations [28] over 10^3^ cells. In both panels, the dashed line indicates the identity *y* = *x* for reference.

**Table 1:**
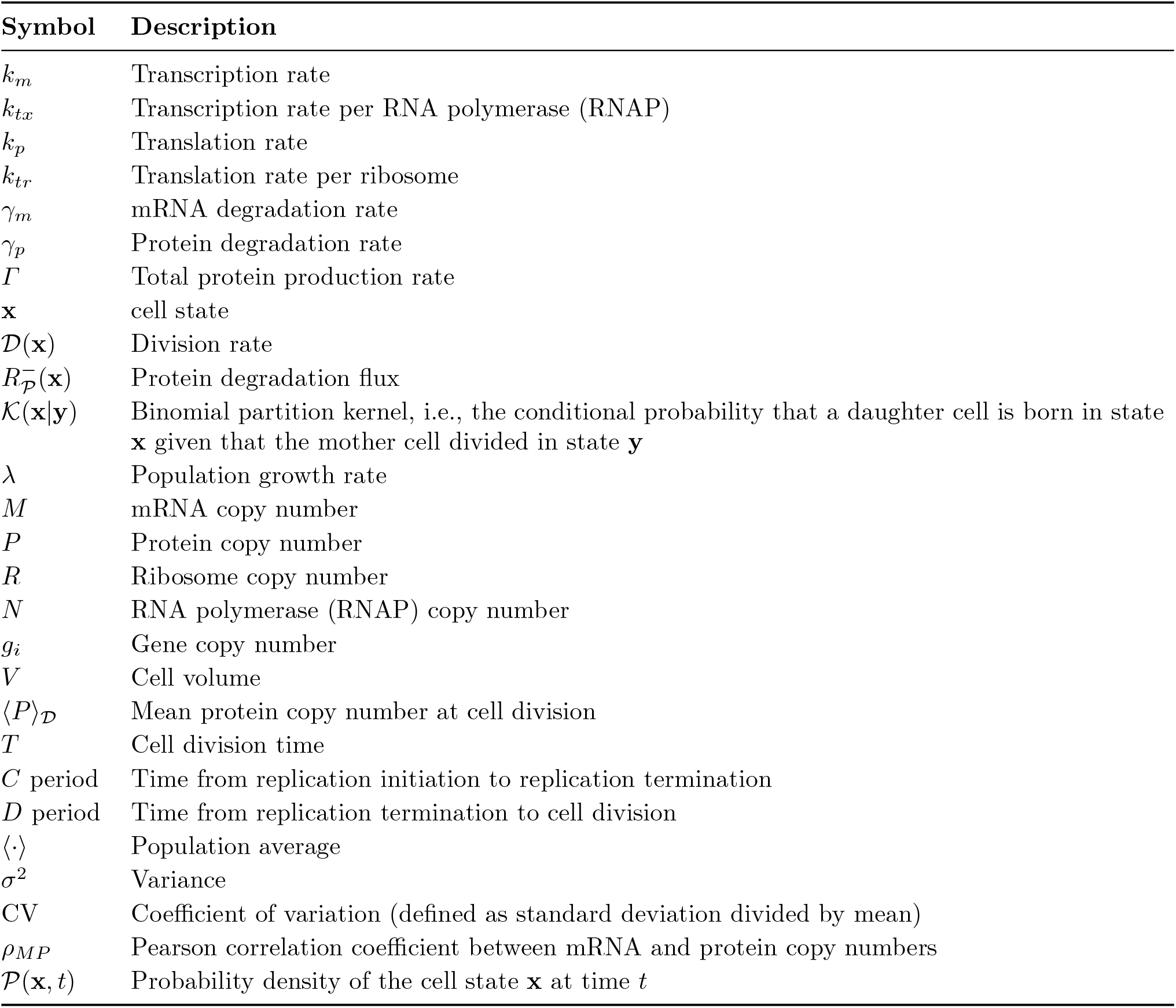
Description of symbols used throughout the manuscript.

## References

[1] M. B. Elowitz, A. J. Levine, E. D. Siggia, and P. S. Swain, Science 297, 1183 (2002).

[2] M. Thattai and A. van Oudenaarden, Proceedings of the National Academy of Sciences of the United States of America 98, 8614 (2001).

[3] P. S. Swain, M. B. Elowitz, and E. D. Siggia, Proceedings of the National Academy of Sciences of the United States of America 99, 12795 (2002).

[4] J. M. Pedraza and A. van Oudenaarden, Science 307, 1965 (2005).

[5] J. Paulsson, Physics of Life Reviews 2, 157 (2005).

[6] M. Kærn, T. C. Elston, W. J. Blake, and J. J. Collins, Nature Reviews Genetics 6, 451 (2005).

[7] I. Golding and A. Amir, Reviews of Modern Physics 96, 041001 (2024).

[8] A. Hilfinger, T. M. Norman, and J. Paulsson, Cell Systems 2, 251 (2016).

[9] A. Hilfinger, T. M. Norman, G. Vinnicombe, and J. Paulsson, Physical Review Letters 116, 058101 (2016).

[10] E. Joly-Smith, M. M. Talpur, P. Allard, F. Papazotos, L. Potvin-Trottier, and A. Hilfinger, eLife 13, RP92497 (2025).

[11] L. A. Sepúlveda, H. Xu, J. Zhang, M. Wang, and I. Golding, Science 351, 1218 (2016).

[12] D. J. Kiviet, P. Nghe, N. Walker, S. Boulineau, V. Sunderlikova, and S. J. Tans, Nature 514, 376 (2014).

[13] P. Thomas, N. Popović, and R. Grima, Proceedings of the National Academy of Sciences 111, 6994 (2014).

[14] M. Soltani, C. A. Vargas-Garcia, D. Antunes, and A. Singh, PLOS Computational Biology 12, e1004972 (2016).

[15] R. Perez-Carrasco, C. Beentjes, and R. Grima, Journal of the Royal Society Interface 17, 20200360 (2020).

[16] C. Jia, A. Singh, and R. Grima, PLOS Computational Biology 18, e1010574 (2022).

[17] Y. Wang, Z. Yu, R. Grima, and Z. Cao, The Journal of Chemical Physics 159 (2023), 10.1063/5.0173742.

[18] J. Lin and A. Amir, Cell Systems 5, 358 (2017).

[19] Y. Hein and F. Jafarpour, Physical Review Research 6, 043006 (2024).

[20] J. D. Little, Operations Research 9, 383 (1961).

[21] A. Novick and L. Szilard, Science 112, 715 (1950).

[22] V. Bryson and W. Szybalski, Science 116, 45 (1952).

[23] P. A. P. Moran, Mathematical Proceedings of the Cambridge Philosophical Society 54, 60 (1958).

[24] A. L. Goldberg and J. F. Dice, Annual Review of Biochemistry 43, 835 (1974).

[25] V. Shahrezaei and P. S. Swain, Proceedings of the National Academy of Sciences of the United States of America 105, 17256 (2008).

[26] K. R. Ghusinga, J. J. Dennehy, and A. Singh, Proceedings of the National Academy of Sciences of the United States of America 114, 693 (2017).

[27] A. Dal Co, M. Cosentino Lagomarsino, M. Caselle, and M. Osella, Nucleic Acids Research 45, 1069 (2017).

[28] D. T. Gillespie, The Journal of Physical Chemistry 81, 2340 (1977).

[29] J. Lin and A. Amir, Nature Communications 9, 4496 (2018).

[30] T. W. Anderson, An Introduction to Multivariate Statistical Analysis, 3rd ed. (Wiley-Interscience, 2003).

[31] J. S. Verdaasdonk, J. Lawrimore, and K. Bloom, in Methods in cell biology, Vol. 123 (Elsevier, 2014) pp. 347–365.

[32] P. Wang, L. Robert, J. Pelletier, W. L. Dang, F. Taddei, A. Wright, and S. Jun, Current Biology 20, 1099 (2010).

[33] H. Kitano, Science 295, 1662 (2002).

[34] T. Hastie, R. Tibshirani, and J. Friedman, The Elements of Statistical Learning: Data Mining, Inference, and Prediction, 2nd ed., Springer Series in Statistics (Springer, New York, NY, 2009).

[35] C. Jarzynski, Physical Review Letters 78, 2690 (1997).

[36] L. Calabrese, L. Ciandrini, and M. Cosentino Lagomarsino, Proceedings of the National Academy of Sciences 121, e2400679121 (2024).

[37] Y. Taniguchi, P. J. Choi, G.-W. Li, H. Chen, M. Babu, J. Hearn, A. Emili, and X. S. Xie, Science 329, 533 (2010).

[38] M. Streit, M. Budiarta, M. Jungblut, and G. Beliu, Biophysical Reports 5 (2025).

[39] A. Schmidt, G. Gao, S. R. Little, A. P. Jalihal, and N. G. Walter, Wiley Interdisciplinary Reviews: RNA 11, e1587 (2020).

[40] J. Reimegård, M. Tarbier, M. Danielsson, J. Schuster, S. Baskaran, S. Panagiotou, N. Dahl, M. R. Friedläander, and C. J. Gallant, Communications biology 4, 624 (2021).

[41] M. Metelev, E. Lundin, I. L. Volkov, A.H. Gynnå, J. Elf, and M. Johansson, Nature communications 13, 1852 (2022).

[42] S. Cooper and C. E. Helmstetter, Journal of Molecular Biology 31, 519 (1968).

[43] W. D. Donachie, Nature 219, 1077 (1968).

[44] A. Amir, Physical Review Letters 112, 208102 (2014).

[45] F. Jafarpour, “Exactly solvable population model with square-root growth noise and cell-size regulation,” (2025), arXiv:2512.05190v1, 2512.05190 [q-bio.PE] .

[46] H. Kempe, A. Schwabe, F. Crémazy, P. J. Verschure, and F. J. Bruggeman, Molecular Biology of the Cell 26, 797 (2015).

[47] R. Ietswaart, S. Rosa, Z. Wu, C. Dean, and M. Howard, Cell Systems 4, 622 (2017).

